# Noncanonical Short-Latency Auditory Pathway Directly Activates Deep Cortical Layers

**DOI:** 10.1101/2025.01.06.631598

**Authors:** Michellee M. Garcia, Amber M Kline, Koun Onodera, Hiroaki Tsukano, Pranathi R. Dandu, Hailey C. Acosta, Michael Kasten, Paul B. Manis, Hiroyuki K. Kato

**Affiliations:** Department of Psychiatry, University of North Carolina at Chapel Hill, Chapel Hill, NC 27599, USA; Neuroscience Center, University of North Carolina at Chapel Hill, Chapel Hill, NC 27599, USA; Department of Otolaryngology/Head and Neck Surgery, University of North Carolina at Chapel Hill, Chapel Hill, NC 27599, USA; Department of Cell Biology and Physiology, University of North Carolina at Chapel Hill, Chapel Hill, NC 27599, USA; Eaton-Peabody Laboratories, Massachusetts Eye and Ear, Boston, MA 02114, USA; Department of Otolaryngology - Head and Neck Surgery, Harvard Medical School, Boston, MA 02114, USA

## Abstract

Auditory processing in the cerebral cortex is considered to begin with thalamocortical inputs to layer 4 (L4) of the primary auditory cortex (A1). In this canonical model, A1 L4 inputs initiate a hierarchical cascade, with higher-order cortices receiving pre-processed information for the slower integration of complex sounds. Here, we identify alternative ascending pathways in mice that bypass A1 and directly reach multiple layers of the secondary auditory cortex (A2), indicating parallel activation of these areas alongside sequential information processing. We found that L6 of both A1 and A2 receive short-latency (<10 ms) sound inputs, comparable in speed to the canonical A1 L4 input but transmitted through higher-order thalamic nuclei. Additionally, A2 L4 is innervated by a caudal subdivision within the traditionally defined primary thalamus, which we now identify as belonging to the non-primary system. Notably, both thalamic regions receive projections from distinct subdivisions of the higher-order inferior colliculus, which in turn are directly innervated by cochlear nucleus neurons. These findings reveal alternative ascending pathways reaching A2 at L4 and L6 via secondary subcortical structures. Thus, higher-order auditory cortex processes both slow, pre-processed information and rapid, direct sensory inputs, enabling parallel and distributed processing of fast sensory information across cortical areas.

## Main Text

Sensory information reaches the cerebral cortex through multiple ascending pathways, with parallel routes existing even within a single sensory modality^1–7^. These pathways are thought to extract and transmit different aspects of the sensory environment, though their specific roles remain largely unclear. Characterizing the unique information each pathway conveys and determining how their inputs are spatially and temporally distributed within the cortex is crucial for understanding how the brain ultimately integrates these inputs to create unified sensory perceptions.

In the auditory system, the cortex receives input through two anatomically distinct pathways: the lemniscal and non-lemniscal pathways^8,9^. The lemniscal pathway originates in the cochlear nucleus, ascends through the central nucleus of the inferior colliculus (CIC) to the ventral division (MGv) of the medial geniculate nucleus (MGN), and terminates in the primary auditory cortex. This pathway is characterized by auditory-specific, short-latency responses with tonotopic organization and is thus considered the primary ascending route. In contrast, the non-lemniscal pathways follow complementary routes, arising from the external and dorsal cortices of the inferior colliculus (ECIC and DCIC), traveling through the dorsal and medial divisions of the MGN (MGd and MGm), and projecting broadly across auditory cortical areas. Neurons in the non-lemniscal pathways typically exhibit longer-latency responses with broader frequency tuning, receive dense cortical feedback, and often respond to non-auditory stimuli^8,10–16^. These distinct characteristics have fostered the view that non-lemniscal pathways serve either as conduits from the primary to secondary auditory cortices or as slower, modulatory routes for integrating multisensory signals, rather than as ascending pathways carrying peripheral sound information^17–24^.

At the thalamocortical level, lemniscal projections from MGv to L4 of the primary cortex have been considered the initial entry point for auditory signals. This “canonical cortical circuit” model^25^ posits that sensory information flows sequentially within the cortical column along the L4→L2/3→L5/6 axis before being transmitted to subcortical structures and higher-order cortices^23,24^ (Fig. 1a). However, emerging findings suggest the presence of ascending sensory inputs with short latencies directly reaching deeper cortical layers^26,27^ and higher-order auditory cortices^28–34^, calling into question whether this strictly hierarchical model fully captures the complexity of thalamocortical processing. While these short-latency inputs have typically been attributed to collaterals of L4-projecting lemniscal thalamocortical axons, an alternative possibility remains unexplored: non-lemniscal pathways, which project broadly across cortical layers and areas^35–38^, might serve as independent, parallel ascending routes that directly carry peripheral sound information to the auditory cortex.

**Fig. 1.**
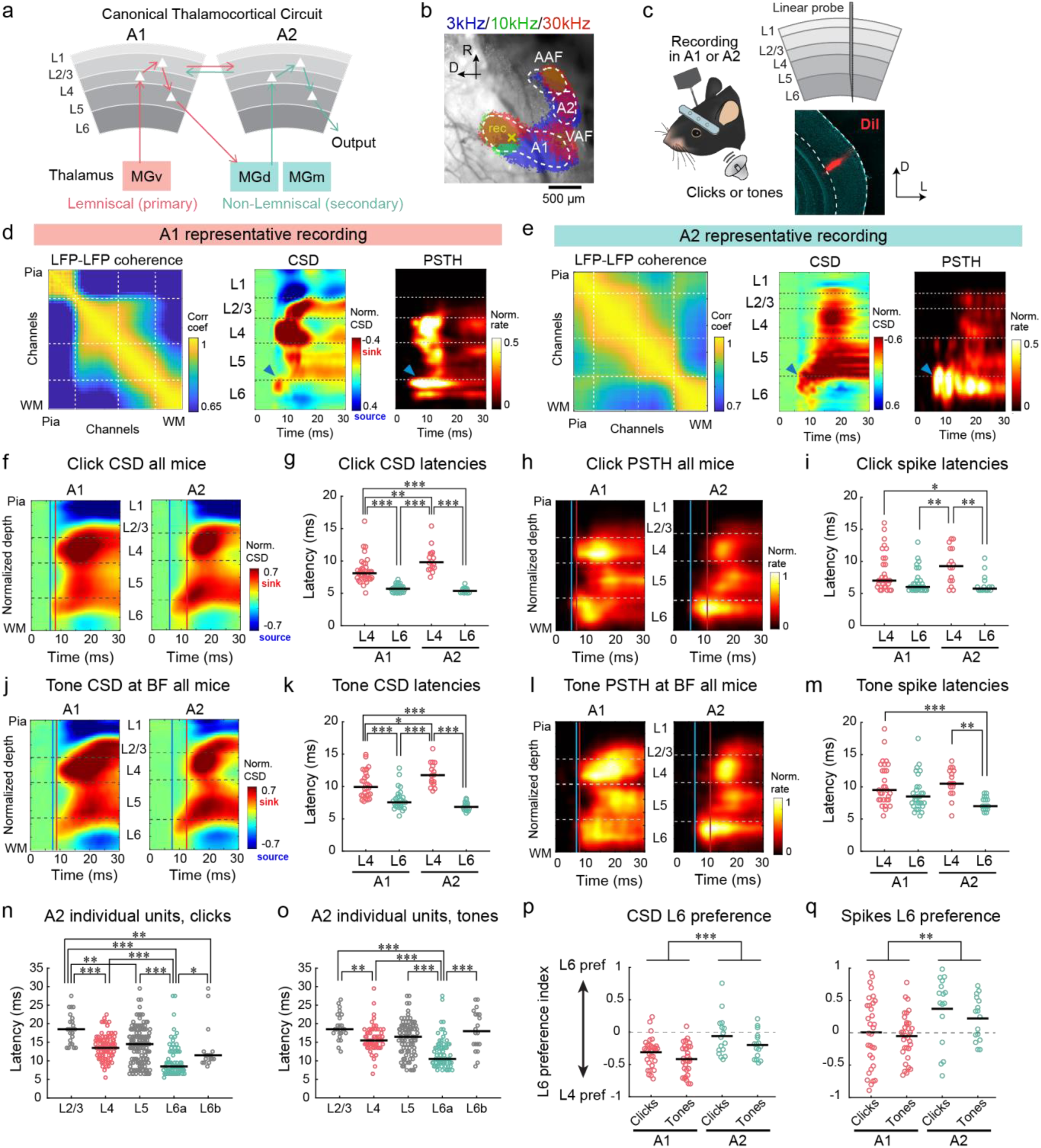
L6 in both A1 and A2 receives short-latency sound inputs. **(a)** Schematic of the canonical thalamocortical circuit. **(b)** Intrinsic signal imaging of pure tone responses superimposed on the cortical surface imaged through the skull. Dotted lines: auto-sorted area borders. Yellow cross: recording site. **(c)** Illustration of linear probe recording across the cortical column. Bottom right: representative probe track in A1 marked by DiI. **(d)** Representative A1 recording data with laminar boundaries. Left: pairwise LFP coherence between channels. Middle: click-triggered current source density (CSD) signals. Right: peristimulus time histogram (PSTH). Blue arrowhead: short-latency L6a response. **(e)** Same as (d), but for A2 recording. **(f)** Click-triggered CSD signals normalized and averaged across all A1 (left) and A2 (right) recordings. Red and blue lines: response onsets in L4 and L6, respectively. **(g)** Summary of CSD response latencies (A1: *n* = 31 mice; A2: *n* = 16 mice; \**p* < 0.05, \*\**p* < 0.01, \*\*\**p* < 0.001, two-way ANOVA followed by Tukey’s HSD test). Black bars: median. **(h)** Same as (f), but for click-triggered PSTHs. **(i)** Summary of spike response latencies. **(j–m)** Same as (f–i), but for BF tone-triggered responses (A1: *n* = 31 mice; A2: *n* = 16 mice). **(n)** First-spike latencies of A2 click responses in individual units by layer. Two-sided Wilcoxon rank sum test with Bonferroni correction. **(o)** First-spike latencies of A2 BF tone responses in individual units by layer. **(p)** L6 preference index of CSD signals for each mouse-area-sound combination (A1 vs. A2: \*\*\**p* = 1.95×10^−5^; click vs. tone: *p*\* = 0.0213; two-way ANOVA). The index is calculated as (L6 response − L4 response) / (L6 response + L4 response). **(q)** Same as (p) but for spike responses (A1 vs. A2: \*\**p* = 0.0011; click vs. tone: *p* = 0.360).

To test this hypothesis and comprehensively delineate parallel ascending pathways, we combined anatomical tracing and electrophysiological recordings to identify the sources of inputs to different layers in both A1 and A2. Our experiments revealed two distinct non-lemniscal ascending pathways that can be traced back to the cochlear nucleus: a short-latency pathway from MGm to L6 of both A1 and A2, and a slower route to A2 L4 from the caudal MGv, which we unexpectedly identified as part of the non-lemniscal system. These findings challenge the traditional model of hierarchical cortical activation by demonstrating that parallel pathways convey distinct streams of auditory information to specific cortical layers and regions, thereby enabling fast and distributed sensory processing.

## Results

### L6 of both A1 and A2 receives short-latency sound inputs with broad frequency tuning

To investigate sound response latencies across cortical layers in A1 and A2, we conducted electrophysiological recordings in awake head-fixed mice. Using intrinsic signal imaging through the intact skull, we first mapped auditory cortical areas by presenting pure tones at three frequencies (3, 10, and 30 kHz; 75 dB SPL; 1 s)^39^ (Fig. 1b). Following this functional mapping, we inserted a linear probe perpendicular to the cortical surface to record simultaneously from all six layers (Fig. 1c). Cortical layer boundaries were determined using a combination of current source density (CSD) signals, spikes, and coherence between local field potentials (LFPs) at individual channels^40^ (see Methods).

When presenting broadband click sounds, we observed distinct response latencies across cortical layers in both CSD signals and multi-unit spikes (Fig. 1d,e). In A1, a short-latency CSD sink consistently appeared in the middle layer, indicating the arrival of fast lemniscal input to L4 (Fig. 1d,f). The L4 sink was significantly slower in A2, consistent with previous studies^41,42^ (Fig. 1e–g, A1 L4 vs. A2 L4: *p* = 0.0032; *n* = 31 and 16 mice for A1 and A2; two-way ANOVA followed by Tukey’s HSD test). Notably, we observed short-latency CSD sinks in L6a of both A1 and A2, which were significantly faster than those in L4 (A1 L4 vs. L6: *p* = 0.0032; A2 L4 vs. L6: *p* = 3.8×10^−9^) but similar between areas (A1 L6 vs. A2 L6: *p* = 0.88). We confirmed these results in multi-unit spike latencies at both population (Fig. 1h,i) and individual unit levels (Fig. 1n and Supplementary Fig. 1a). The temporal difference was particularly striking in A2, where L6 input preceded L4 by approximately 5 ms. This timing difference cannot be explained by axon conduction delay alone, which is estimated to be 1.2 ms based on a 0.4 mm travel distance between L6 and L4, assuming 0.33 m/s conduction velocity^43^. This observation argues against the possibility that the fast input in deep layers originates from collaterals of L4-targeting lemniscal axons. No difference was found in latencies between L6 units with different spike waveforms (Supplementary Fig. 1c–e). These findings demonstrate short-latency sound responses in A2 and suggest the existence of alternative pathways delivering fast sound inputs to the deep layers of both A1 and A2.

To determine whether the short-latency inputs to L6 were specific to broadband clicks or generalized across different auditory stimuli, we presented pure tones with a wide range of frequencies (4–64 kHz).

At the best frequency (BF) for each recording site, pure tones evoked short-latency responses in L6 of both A1 and A2, as evidenced by both CSD signals and spike responses (Fig. 1j–m,o; Supplementary Fig. 1b). Response latencies to pure tones were approximately 2 ms slower than those to clicks due to the tones’ linear rise function over 5 ms; nonetheless, the L6 responses in both A1 and A2 consistently preceded the L4 inputs. Therefore, rapid sound input to L6 represents a general feature of auditory processing rather than a stimulus-specific phenomenon.

We quantified the relative magnitudes of L4 and L6 inputs by calculating the L6 preference index for each area and sound type (see Methods). This analysis revealed that A2 had significantly higher index values than A1 for both CSD and spike responses (A1 vs. A2, CSD: *p* = 1.95×10^−5^; spikes: *p* = 0.0011; *n* = 20 and 9 mice for A1 and A2; two-way ANOVA), indicating a stronger contribution of L6 input in A2 (Fig. 1p,q). Additionally, click sounds showed higher L6 preference than tones (clicks vs. tones, CSD: *p* = 0.0213; spikes: *p* = 0.360), suggesting distinct sound preferences between L4 and L6 inputs.

To further characterize the distinct tuning properties of L4 and L6 inputs and their receiver neurons, we analyzed the frequency tuning broadness of both CSD and spikes. We quantified response magnitudes within narrow time windows around response onsets (5–15 ms after tone onsets for A1 L4, A1 L6, and A2 L6; 10–20 ms after tone onsets for A2 L4) to isolate pathway-specific inputs and minimize contributions from across-layer propagation. As shown in a representative A1 recording, the onset response in A1 L4 exhibits sharp frequency tuning, as expected for lemniscal input (Fig. 2a). In contrast, the A1 L6 input in the same recordings exhibited significantly broader frequency tuning, as measured by full width at half maximum (FWHM) of both CSD and spikes (A1 L4 vs. A1 L6, CSD: *p* = 3.73×10^−5^; spikes: *p* = 1.83×10^−5^; two-way ANOVA followed by Tukey’s HSD test) (Fig. 2a–e). In A2, both L4 and L6 exhibited broad frequency tuning comparable to that observed in A1 L6 (A2 L4 vs. A2 L6, CSD: *p* = 0.999; spikes: *p* = 0.53; A1 L6 vs. A2 L6, CSD: *p* = 0.872; spikes: *p* = 0.968). Similar results were obtained when measuring the tuning of regular-spiking units (Supplementary Fig. 2). This difference in tuning properties suggests that L4 and L6 inputs in A1 either have different across-frequency convergence or convey distinct sound information and likely originate from separate pathways. Regardless of the exact circuit, these findings demonstrate that inputs to L6 in both A1 and A2 share similar characteristics: short-latency responses and broad frequency tuning.

**Fig. 2.**
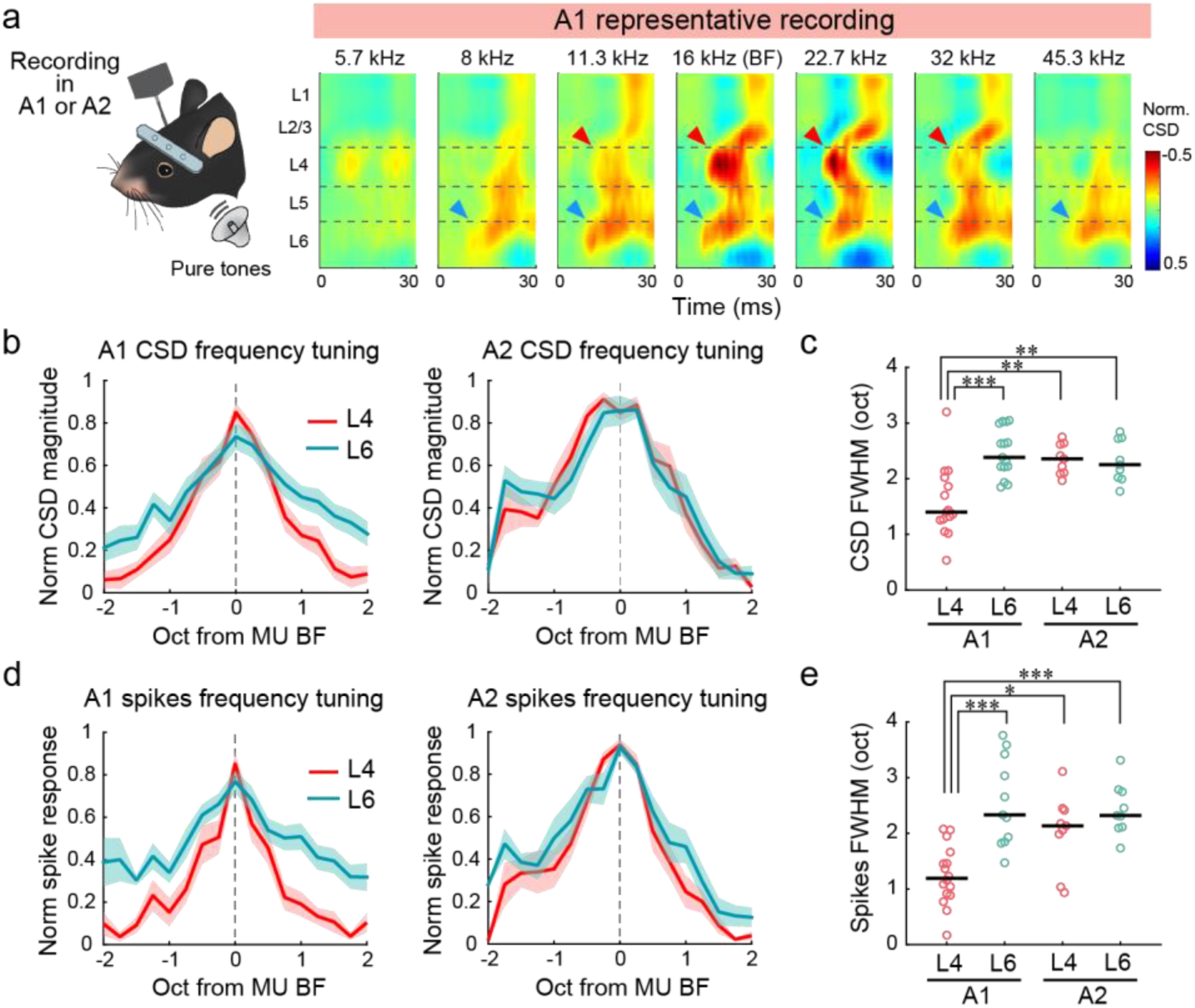
L6 input shows broader frequency tuning than L4 input. **(a)** Representative A1 recording showing CSD signals across tone frequencies. Red and blue arrowheads: responses in L4 and L6, respectively. **(b)** Normalized frequency tuning curves of L4 and L6 CSD sinks in A1 (left) and A2 (right), centered at the BF of the recording site (A1: *n* = 20 mice; A2: *n* = 9 mice). Lines: mean. Shading: SEM. **(c)** Summary of full-width half-maximum (FWHM) of tuning curves (\**p* < 0.05, \*\**p* < 0.01, \*\*\**p* < 0.001, two-way ANOVA followed by Tukey’s HSD test). Black bars: median. **(d–e)** Same as (b**–**c) but for spike responses.

### Distinct thalamic origins for parallel ascending pathways onto different cortical layers

A previous tracing study identified projections from the caudal subdivision of MGv (cMGv) to A2^34^. Therefore, a parsimonious explanation for the short-latency responses in A2 L6 may be its direct innervation by the lemniscal MGv. To test this hypothesis and determine the thalamic origins of parallel pathways to L4 and L6 in both A1 and A2, we conducted anatomical tracing experiments. Following functional mapping of auditory cortical areas using intrinsic signal imaging, we targeted retrograde tracer injections to L4 or L6 of these identified regions (Fig. 3a). Injections into A1 L4 resulted in retrograde labeling predominantly in the rostral subdivision of MGv (rMGv), consistent with the canonical cortical circuit model (Fig. 3b,d). Similar injections in A2 L4 led to predominant labeling in cMGv, confirming previous findings (Fig. 3c,e). Additionally, A2 L4 received stronger projections from MGd compared to A1, aligning with A2’s role as a higher-order cortex. The rostrocaudal distribution of inputs revealed an approximate boundary at −3.30 mm from bregma, dividing MGv into subdivisions projecting to A1 L4 and A2 L4 (Fig. 3f).

**Fig. 3.**
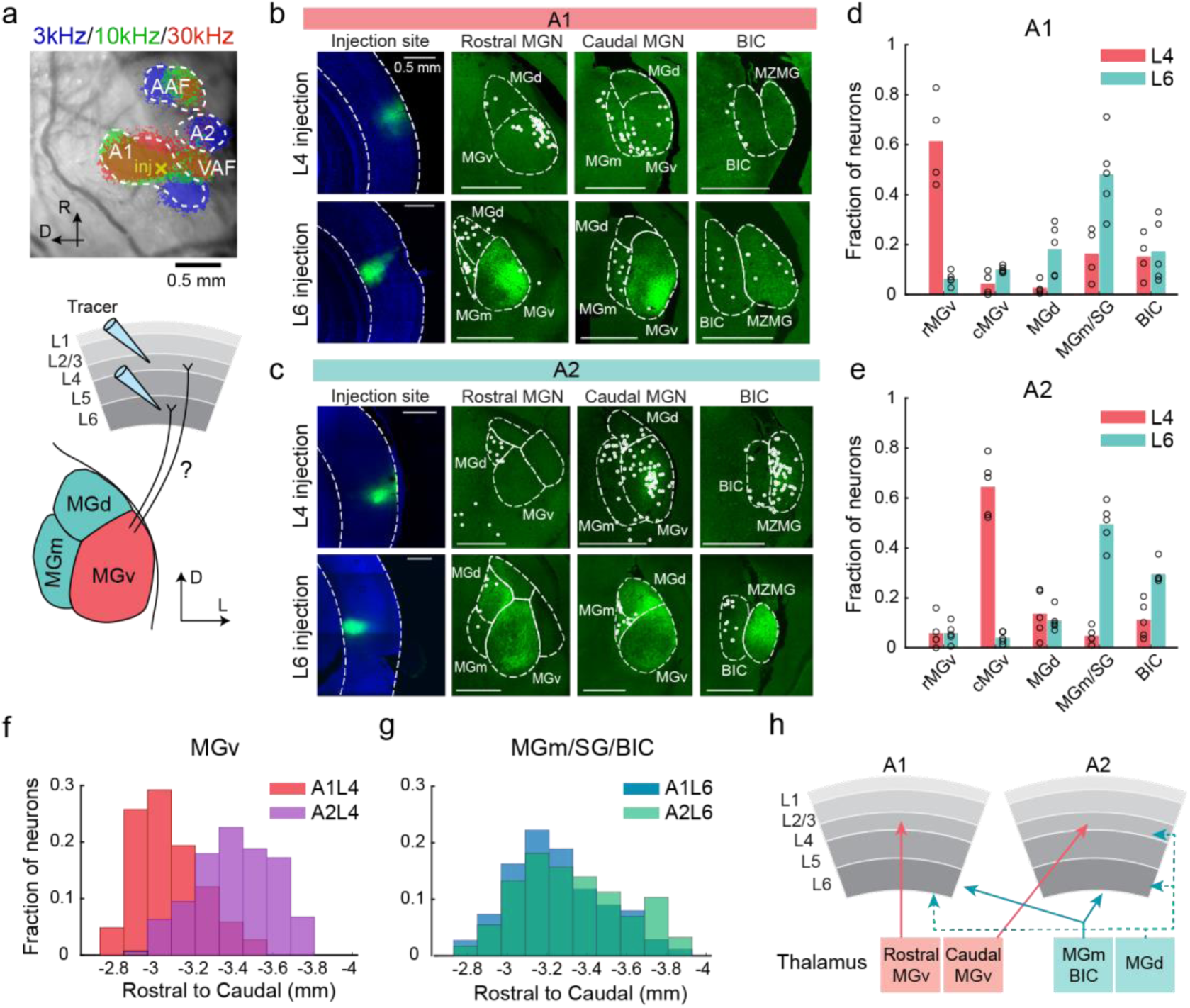
Distinct thalamic origins for parallel ascending pathways to cortical layers. **(a)** Top: intrinsic signal imaging of pure tone responses superimposed on the cortical surface imaged through the skull. Yellow cross: injection site. Bottom: diagram of layer-targeted retrograde tracing. **(b)** Representative tracing results of CTB488 injections in A1 L4 (top) and L6 (bottom). White circles: input cells shown in coronal sections at rostral MGN, caudal MGN, and BIC levels. Note that diffuse MGv signals in L6 injections represent anterogradely labeled L6 axons. MZMG: marginal zone of the medial geniculate. **(c)** Same as (b), but for A2 injections. **(d)** Fraction of input cells across MGN divisions for A1 L4 and L6 injections (L4: *n* = 4 mice; L6: *n* = 5 mice). For each injection, the fraction of labeled neurons in each input region was calculated relative to the total MGN input. Bar heights: mean. **(e)** Same as (d), but for A2 injections (L4: *n* = 4 mice; L6: *n* = 5 mice). **(f)** Rostrocaudal distribution of MGv input cells to L4 in A1 (red) and A2 (purple). **(g)** Rostrocaudal distribution of MGm/SG/BIC input cells to L6 in A1 (blue) and A2 (green). **(h)** Revised thalamocortical connectivity schematic.

Surprisingly, L6 of both A1 and A2, despite exhibiting short-latency responses, received minimal input from MGv. Instead, the majority of projections to L6 originated from medial structures: MGm, the suprageniculate nucleus (SG), and the nucleus of the brachium of the inferior colliculus (BIC) (Fig. 3b– e). Due to the small number of SG-labeled neurons and unclear borders between structures, we combined SG with MGm for quantification purposes. The rostrocaudal distributions of A1 L6- and A2 L6-projecting neurons in these medial structures largely overlapped, with A2-projecting neurons displaying a slight caudal bias (Fig. 3g). The identification of BIC as a source of cortical projections was unexpected, as BIC has traditionally not been considered part of the thalamic nuclei. However, our results suggest that the rostral subdivision of BIC (rBIC; from −3.5 to −4.1 mm caudal from bregma) associates with thalamic nuclei and provides direct projections to the cortex (see Discussion and Supplementary Fig. 3 for more detailed analyses).

Together, these findings demonstrate that while L4 of both A1 and A2 receives MGv inputs, L6 exhibits a complementary pattern of inputs from the medial MGN structures. These distinct input patterns, along with electrophysiological data (Figures 1 and 2), further argue against the hypothesis that short-latency inputs to deep layers arise from collaterals of L4-targeting axons, requiring a revision of the thalamocortical circuit model (Fig. 3h).

### MGm and BIC subpopulations transmit sound information with short latencies

Our anatomical results revealed distinct thalamic origins for parallel pathways to L4 and L6, raising two key questions. First, do MGm and rBIC respond rapidly enough to account for the observed short-latency inputs to L6? Second, given the long-latency responses in A2 L4, does cMGv convey slower sound responses despite its presumed lemniscal identity? To address these questions, we conducted linear probe recordings in 26 mice, sampling all MGN divisions and rBIC (Fig. 4a). Probe trajectories were reconstructed and mapped to MGN divisions using the Paxinos Brain Atlas^44^ (Fig. 4b,c; see Methods). We chose the Paxinos Brain Atlas over the Allen Brain Atlas for MGN mapping, as the former provides a more precise delineation of MGN divisions. In recordings that sampled both MGv and MGm simultaneously, we observed a clear transition in neural activity patterns at the MGv-MGm border (Supplementary Fig. 4), validating that our spatial mapping captures functional divisions of MGN.

**Fig. 4.**
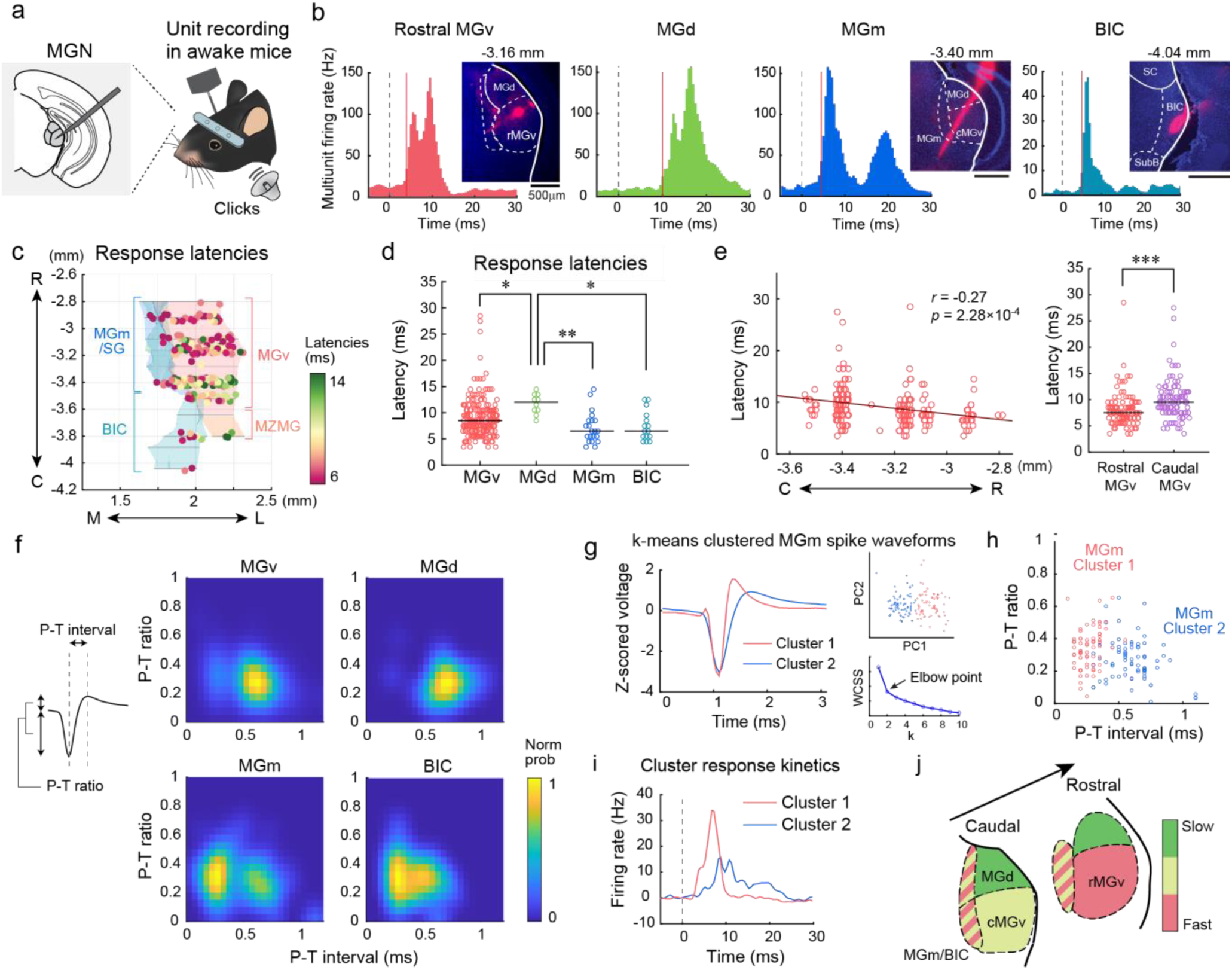
Neural subpopulations in MGm and BIC transmit short-latency sound information. **(a)** MGN linear probe recording illustration. **(b)** Representative click-triggered PSTHs for channels in rMGv, MGd, MGm, and BIC. Red lines: first-spike latencies. Adjacent histology images: corresponding coronal brain sections with DiI-marked probe tracks. **(c)** Click response latencies at reconstructed recording sites across MGN divisions. Dorsal view of 3-D reconstructions from 26 mice. MGd excluded to visualize the transition between cMGv and MGm/BIC. **(d)** First-spike latencies of click responses in individual units for each MGN division (\**p* < 0.05, \*\**p* < 0.01, \*\*\**p* < 0.001, two-sided Wilcoxon rank sum test with Bonferroni correction). Black lines: median. **(e)** Left: distribution of click response latencies in MGv along the rostrocaudal axis (Pearson’s correlation; two-sided t-test). Red line: linear fit. Right: click response latencies in rMGv vs. cMGv (\*\*\**p* = 1.86×10^−5^; two-sided Wilcoxon rank sum test). **(f)** Left: P-T interval and P-T ratio example. Right: two-dimensional heat maps showing the probability distribution of P-T intervals (x-axis) and P-T ratio (y-axis) for each MGN division. **(g–i)** k-means clustering of MGm spike waveforms. **(g)** Left: averaged spike waveforms for units in Cluster 1 (red) and Cluster 2 (blue). Top right: two clusters of MGm spikes in PC space. Bottom right: elbow point analysis (optimal cluster number = 2). WCSS: within-cluster sum of squares. **(h)** Distributions of P-T interval and P-T ratio for individual MGm clusters. **(i)** Click-triggered PSTHs for individual MGm clusters. Cluster 1 (short P-T intervals) shows short-latency responses. **(j)** Illustration of response latencies across MGN divisions. MGm shows heterogeneous latencies across subpopulations.

We measured click response latencies for multi-unit channels with significant sound responses (see Methods). As expected, MGv exhibited short latencies (9.0 ± 0.3 ms), while MGd had longer latencies (11.7 ± 0.6 ms) (Fig. 4b,d). Further analysis revealed a rostrocaudal gradient in MGv response latencies, with cMGv showing significantly longer latencies than rMGv^45^ (*p* = 1.86×10^−5^; Wilcoxon rank sum test) (Fig. 4e). The functional difference between rMGv and cMGv is likely even greater than our measurements suggest, as our simple border plane of −3.30 mm posterior to bregma does not fully capture the real shape of the boundary between these subdivisions. Nevertheless, this difference in response timings explains why A2 L4, which receives projections from cMGv, exhibits delayed responses. MGm and rBIC showed latencies comparable to or faster than MGv (MGm: 7.2 ± 0.6 ms; rBIC: 7.5 ± 0.7 ms), consistent with previous reports of heterogeneous MGm latencies, including those similar to MGv^8,45–47^. Notably, the earliest spikes in MGm and rBIC neurons preceded those in cortical L6, satisfying the necessary causal requirement for these cells to serve as sources of short-latency responses in L6. Examining the 5th percentile latencies across all units revealed a temporal hierarchy: in the thalamus, MGm (3.5 ms), rBIC (4.5 ms), and MGv (4.5 ms) showed the shortest latencies, followed by MGd (8.5 ms). In the cortex, A1 L4, A1 L6, and A2 L6 responded earliest (6.5 ms each), while A2 L4 responded later (8.5 ms).

We next analyzed spike waveform parameters and identified two distinct MGm subpopulations, separated by peak-trough (P-T) intervals around 0.4 ms (Fig. 4f; Supplementary Fig. 5). Applying k-means clustering to spike waveforms confirmed that P-T intervals were the primary distinguishing factor between these clusters (Fig. 4g,h). Averaged peri-stimulus time histograms (PSTHs) revealed that units with short P-T intervals showed short-latency responses, while those with broader spikes had slower, sustained responses (Fig. 4i). This heterogeneity within MGm neurons suggests different functional roles: narrow-spike neurons mediate short-latency signals to L6 of the auditory cortex, whereas broad-spike neurons process slower inputs, potentially including cortical feedback. Although rBIC also showed two clusters, they were less distinct, and both exhibited short-latency responses (Supplementary Fig. 5). In contrast, MGv and MGd primarily featured spikes with long P-T intervals (>0.4 ms). This finding confirms that the short-latency responses observed in MGm and rBIC, marked by narrow P-T intervals, were not due to contamination from MGv neurons. Together, these findings support our anatomical tracing results, revealing three parallel pathways with distinct kinetics: a fast pathway from rMGv to A1 L4, a slower pathway from cMGv to A2 L4, and another fast pathway from MGm/rBIC to L6 in both A1 and A2 (Fig. 3h and 4j).

### Non-lemniscal origins of ascending pathways to cMGv, MGm, and rBIC

To identify the sources of input to MGN subdivisions, we conducted retrograde tracing experiments. The MGN’s small size and deep location present a challenge for stereotaxic targeting of specific subdivisions without cross-subdivision contamination. To address this, we aspirated the overlying cortex and hippocampus to directly visualize the MGN for targeted tracer injections (Fig. 5a). This approach enabled precise, subdivision-restricted delivery of retrograde tracers to rMGv, cMGv, and MGd (Fig. 5b, Supplementary Fig. 6).

**Fig. 5.**
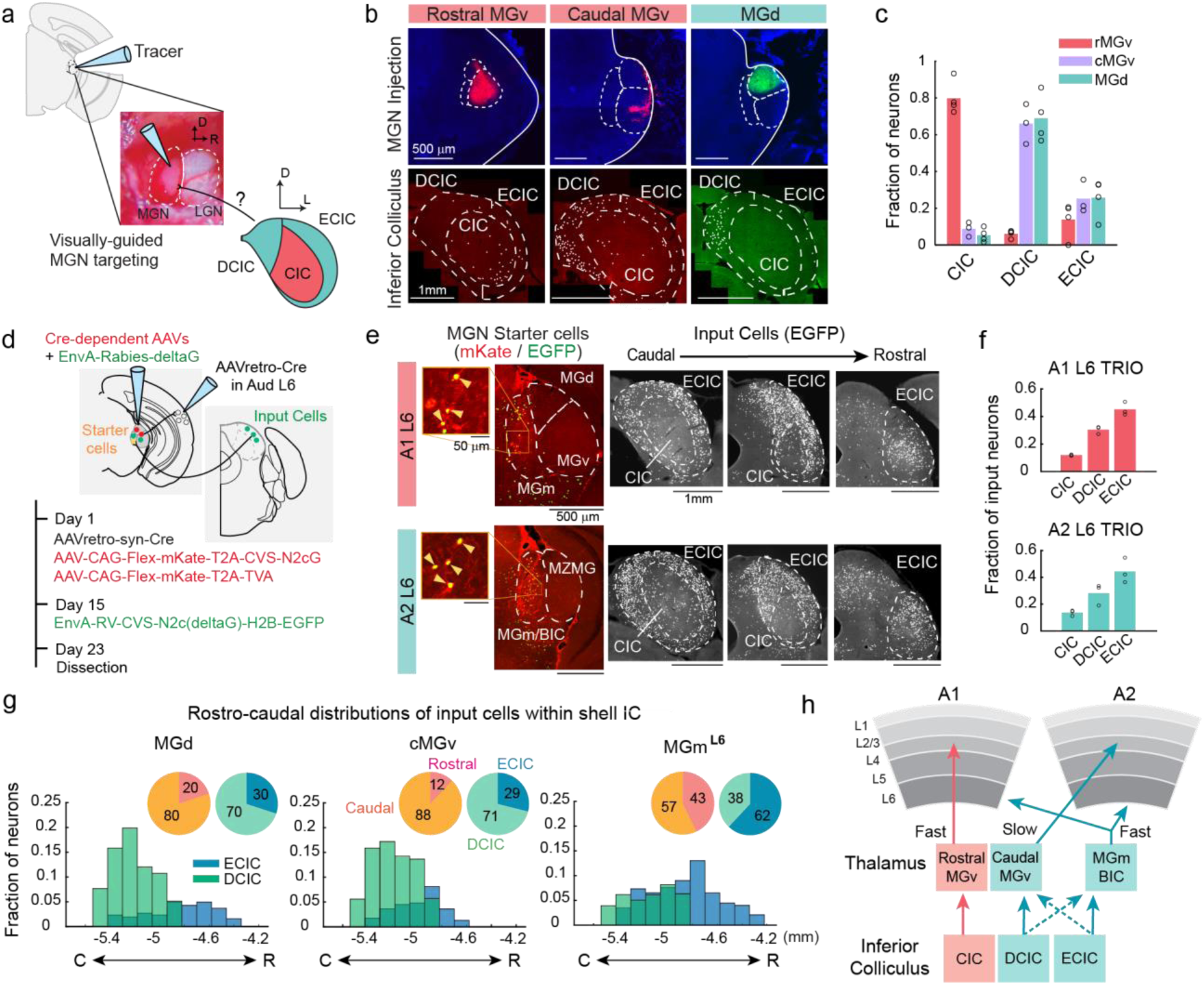
Non-lemniscal origins of ascending pathways to cMGv, MGm, and rBIC. **(a)** Illustration of retrograde tracer injection into the exposed MGN. Middle: Lateral view of exposed MGN and lateral geniculate nucleus (LGN) after cortex and hippocampus aspiration. **(b)** Representative tracing results of CTB594 (rMGv), retrobeads (cMGv), and CTB488 (MGd) injections. Top: coronal MGN sections at injection sites. Bottom: caudal inferior colliculus sections with input cells (white dots). **(c)** Input cell distribution across inferior colliculus divisions for rMGv, cMGv, and MGd injections (rMGv: *n* = 4 mice; cMGv: *n* = 3 mice; MGd: *n* = 4 mice). For each injection, the fraction of labeled neurons in each input region was calculated relative to the total inferior colliculus input. Bar heights: mean. **(d)** Illustration of disynaptic rabies tracing (TRIO method). **(e)** Representative TRIO tracing results from A1 L6 (top) and A2 L6 (bottom). Left: starter cells in MGm/BIC expressing mKate- and EGFP. Insets: magnified views of starter cells (arrowheads). Right: coronal brain sections showing EGFP-expressing input cells at caudal, middle, and rostral inferior colliculus levels. **(f)** Input cell distribution across inferior colliculus divisions for A1 L6 TRIO (top) and A2 L6 TRIO (bottom) (A1: *n* = 3 mice; A2: *n* = 3 mice). **(g)** Rostrocaudal distribution of input cells to MGd (left), cMGv (middle), and MGm^L6^ (right), showing fractions of total counts within the inferior colliculus shell. Blue and green bars: mean of ECIC and DCIC, respectively. Inset pie charts: fraction of rostral (pink) vs. caudal (orange) shell, and ECIC (blue) vs. DCIC (green). **(h)** Revised schematic of ascending pathways through inferior colliculus, thalamus, and cortex. cMGv is classified as part of the non-lemniscal pathway.

Retrograde tracing revealed distinct input patterns for each MGN subdivision. As expected for the canonical lemniscal pathway, injections in rMGv predominantly labeled neurons in the CIC (Fig. 5b,c). In contrast, MGd injections labeled neurons in the shell of the inferior colliculus (DCIC and ECIC), confirming that it receives non-lemniscal input. Notably, cMGv injections resulted in a labeling pattern closely resembling MGd rather than rMGv, with predominant inputs from DCIC and ECIC but minimal labeling in CIC. Therefore, despite the traditional classification of MGv as a homogeneous lemniscal structure, the cMGv is more accurately described as part of the non-lemniscal pathway. This structural organization provides a basis for the long-latency sound responses observed in cMGv and its target, A2 L4.

Our electrophysiological data revealed heterogeneous spike shapes and response latencies in MGm and rBIC neurons. Therefore, to investigate the origin of short-latency responses conveyed to cortical L6, we employed disynaptic retrograde tracing using rabies viruses (TRIO method; Fig. 5d). After mapping cortical areas with intrinsic signal imaging, we injected AAVretro-Cre into either A1 L6 or A2 L6. During the same surgery, we stereotaxically targeted AAV-Flex-mKate-T2A-CVS-N2cG and AAV-Flex-mKate-T2A-TVA to the MGN. Two weeks later, EnvA-Rabies-CVS-N2c(ΔG)-H2B-EGFP was injected at the same MGN location to infect TVA-expressing neurons. This approach successfully labeled mKate- and EGFP-coexpressing “starter” neurons in MGm and rBIC (Fig. 5e). Analysis of EGFP-expressing input cells showed that the presynaptic partners of these MGm/rBIC starter neurons were predominantly located in ECIC and DCIC, but minimally in CIC (Fig. 5e,f; Supplementary Fig. 7). We found only very sparse input cells in the contralateral cochlear nucleus (0.3 ± 0.1% of counts in the inferior colliculus), making it unlikely that the sparse direct projections which bypass the inferior colliculus^48,49^ account for the robust short-latency responses observed in MGm and BIC. Together, these findings demonstrate that L6-projecting MGm/BIC neurons (hereafter referred to as MGm^L6^ neurons) receive most of their input from the non-lemniscal inferior colliculus, despite their role in conveying short-latency sound information. Note that the TRIO strategy was not feasible for tracing the disynaptic source for A2 L4 due to AAVretro’s tropism against infecting MGv neurons.

These findings demonstrate that both the fast pathway innervating MGm^L6^ neurons and the slower pathways innervating cMGv and MGd originate from non-lemniscal regions of the inferior colliculus. Further analysis of the spatial distribution of input cells revealed distinct origins for these parallel pathways. While MGd and cMGv received predominant inputs from DCIC, inputs to MGm^L6^ were primarily localized in ECIC (Fig. 5g). Additionally, we observed a stronger rostral bias in the input cells to MGm^L6^ neurons: 43% of inputs to MGm^L6^ neurons were located rostral to −4.90 mm (the mid-point of the inferior colliculus), compared to only 12% and 20% of input cells for cMGv and MGd, respectively.

In summary, our results reveal an unexpected organization of parallel ascending pathways to the auditory thalamus. While rMGv receives fast input via the canonical lemniscal pathway, cMGv is associated with the non-lemniscal pathway and receives slower inputs from DCIC. Furthermore, MGm and BIC relay short-latency sound information to cortical L6 through inputs from the rostral ECIC, constituting a distinct non-lemniscal pathway (Fig. 5h).

### Rostral ECIC transmits sound information with short latencies

Our anatomical tracing suggested that MGm^L6^ receives short-latency sound information from the non-lemniscal divisions of the inferior colliculus. To test this hypothesis, we examined sound response latencies across inferior colliculus divisions. We performed linear probe recordings in 13 mice and reconstructed probe trajectories using the Allen Common Coordinate Framework (CCF; Fig. 6a, see Methods). We specifically chose the CCF, which defines the anterior CIC boundary approximately 0.4 mm rostral to that in the Paxinos Brain Atlas, to ensure conservative identification of the rostral ECIC.

**Fig. 6.**
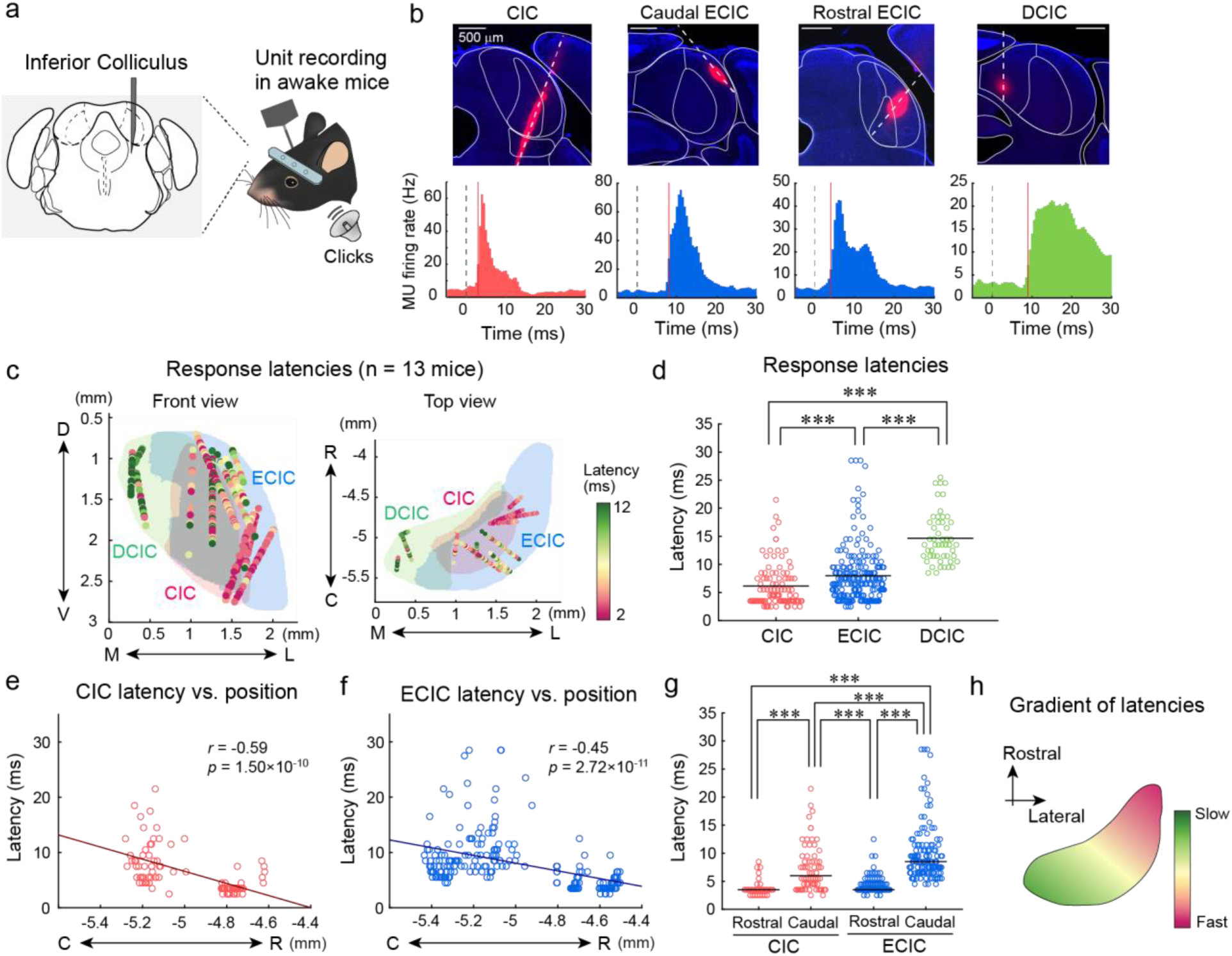
Rostral ECIC transmits sound information with short latencies. **(a)** Inferior colliculus linear probe recording illustration. **(b)** Representative click-triggered PSTHs for channels in CIC, caudal ECIC, rostral ECIC, and DCIC. Red lines: first-spike latencies. Top: coronal sections with DiI-marked probe tracks. **(c)** Click response latencies at reconstructed recording sites across inferior colliculus divisions. Front view (left) and top view (right) of 3-D reconstructions from 13 mice. **(d)** First-spike latencies of click responses in individual units for each inferior colliculus division (CIC: *n* = 7 mice; ECIC: *n* = 10 mice; DCIC: *n* = 4 mice). \*\*\**p* < 0.001, two-sided Wilcoxon rank sum test with Bonferroni correction. Black bars: median. **(e)** Distribution of click response latencies in CIC along the rostrocaudal axis. Red line: linear fit (Pearson’s correlation; two-sided t-test). **(f)** Distribution of click response latencies in ECIC along the rostrocaudal axis. Blue line: linear fit. **(g)** First-spike latencies of click responses in individual units in rostral vs. caudal subdivisions of CIC and ECIC. Two-way ANOVA followed by Tukey’s HSD test. **(h)** Schematic showing a gradient of response latencies along the caudomedial-to-rostrolateral axis.

Click sound responses revealed distinct temporal patterns across divisions of the inferior colliculus. CIC exhibited expected short-latency responses for the lemniscal division (6.2 ± 0.4 ms; 5th percentile: 2.5 ms) (Fig. 6b-d). DCIC showed long-latency responses (14.7 ± 0.6 ms; 5th percentile: 9.5 ms), consistent with the slow responses observed in its targets, MGd and cMGv (Fig. 4d,e). ECIC, as a whole, displayed intermediate response latencies, slightly slower than CIC (8.0 ± 0.3 ms; 5th percentile: 3.5 ms). However, detailed analysis revealed pronounced spatial heterogeneity in response latencies within both CIC and ECIC. We identified a clear rostrocaudal gradient in both divisions, with rostral subdivisions displaying shorter latencies than their caudal counterparts (Fig. 6e-g). Using a rostrocaudal boundary at -4.90 mm, we found significantly shorter latencies in rostral compared to caudal subdivisions. In CIC, the rostral subdivision showed latencies of 3.8 ± 0.3 ms (5th percentile: 2.5 ms) compared to 7.4 ± 0.5 ms in the caudal subdivision (5th percentile: 3.5 ms; *p* = 1.1×10^−4^; two-way ANOVA followed by Tukey’s HSD test). Similarly, in ECIC, the rostral subdivision showed latencies of 4.4 ± 0.2 ms (5th percentile: 2.75 ms) compared to 10.0 ± 0.5 ms in the caudal subdivision (5th percentile: 5.5 ms; *p* = 3.8×10^−9^). Our results demonstrate that both lemniscal and non-lemniscal inferior colliculus display gradients of response latencies along the rostrocaudal axis (Fig. 6h, Supplementary Fig. 8). Notably, rostral ECIC exhibited faster sound responses than caudal CIC (*p* = 6.4×10^−5^), challenging the traditional dichotomy of fast lemniscal CIC and slower non-lemniscal ECIC. This organization explains how the rostral ECIC, which innervates MGm^L6^, can support rapid sound information processing within the non-lemniscal pathway.

### Direct projections from the cochlear nucleus to non-lemniscal inferior colliculus

Finally, to examine whether non-lemniscal divisions of the inferior colliculus receive direct ascending input from the cochlear nucleus, we conducted further retrograde tracing (Fig. 7a). We injected retrograde tracers into four inferior colliculus subdivisions: CIC, DCIC, caudal ECIC, and rostral ECIC. We consistently observed retrogradely labeled cells in the anterior ventral (AVCN), posterior ventral (PVCN), and dorsal cochlear nucleus (DCN), demonstrating direct ascending projections from the cochlear nucleus to all inferior colliculus subdivisions^50–53^ (Fig. 7b,c). Consistent with a recent study showing more CIC-restricted projections from AVCN compared to DCN neurons^51^, retrograde labeling from CIC predominantly labeled AVCN. In contrast, injections in DCIC and caudal ECIC resulted in the predominant labeling of DCN. While we also observed input cells in AVCN and PVCN, these cells tended to be located near the medial, dorsal, and lateral margins of these regions (see caudal ECIC injection in Fig. 7b). Injections in the rostral ECIC resulted in a higher fraction of input cells in AVCN compared to the caudal ECIC, which may underlie the short-latency sound responses in this area. These direct ascending projections provide substrates for the short-latency sound responses observed in non-lemniscal pathways, including the ECIC, MGm, BIC, and L6 of the auditory cortices (Fig. 7d).

**Fig. 7.**
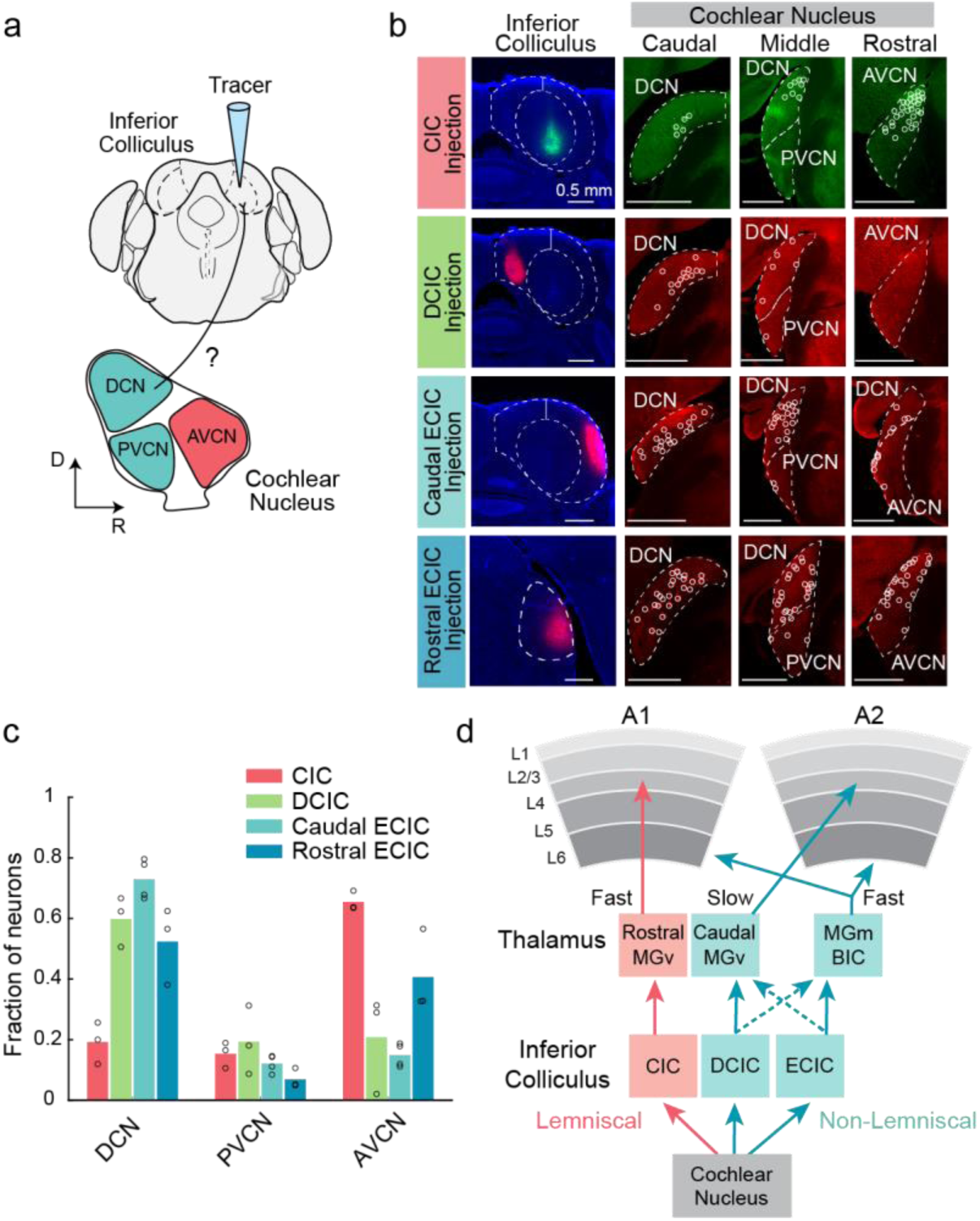
Direct projections from the cochlear nucleus to non-lemniscal inferior colliculus. **(a)** Illustration of retrograde tracing from inferior colliculus divisions. **(b)** Representative retrograde tracing results of tracer injections in CIC, DCIC, caudal ECIC, and rostral ECIC. Left: coronal inferior colliculus sections at injection sites. Right: input cells (white circles) in cochlear nucleus at DCN, PVCN, and AVCN levels. **(c)** Input cell distribution across cochlear nucleus divisions for CIC, DCIC, and ECIC injections (CIC: *n* = 3 mice; DCIC: *n* = 3 mice; caudal ECIC: *n* = 4 mice; rostral ECIC: *n* = 3 mice). For each injection, the fraction of labeled neurons in each input region was calculated relative to the total cochlear nucleus input. Bar heights: mean. **(d)** Revised schematic of parallel ascending pathways from the cochlear nucleus through the inferior colliculus and thalamus to distinct cortical layers.

## Discussion

### Parallel ascending pathways reaching the auditory cortex

Our study revealed two non-lemniscal ascending pathways to the auditory cortex that operate alongside the canonical lemniscal input to A1 L4. First, we identified a neural subpopulation in MGm and BIC that delivers rapid sound information to L6 in both A1 and A2, preceding even the lemniscal input. Second, we demonstrated that the caudal subdivision of MGv, traditionally considered part of the lemniscal pathway, receives predominantly non-lemniscal input and transmits slower sound information to A2 L4. These findings reveal the convergence of at least three parallel ascending pathways that convey sound signals to the auditory cortex. The auditory cortical circuit thus constructs a unified representation of the acoustic environment by integrating these parallel inputs across different layers and areas.

Previous electrophysiological recordings in humans and primates have hinted at the existence of parallel ascending pathways reaching the higher-order auditory cortices^28–33^. For instance, the human superior temporal gyrus (STG), a higher-order cortex essential for speech perception, exhibits sound response latencies comparable to those in the primary Heschl’s gyrus^28^. While these findings indicated the presence of alternative pathways that bypass the primary cortex, their anatomical origins remained unclear. Our study now identifies potential neural substrates for these parallel ascending pathways to higher-order cortical areas that may be conserved across species.

Earlier tracing studies identified parallel inputs from different MGv subdivisions to multiple cortical regions, including A1, A2, anterior auditory field (AAF), ventral auditory field (VAF), and insular auditory field (IAF)^2,34,54–57^. These observations led some researchers to propose reclassifying A2 as a primary rather than higher-order cortex^34,58^. However, our findings challenge the conventional view of MGv as a uniform lemniscal structure. We demonstrate that cMGv receives input predominantly from the shell of the inferior colliculus, particularly DCIC, rather than the lemniscal CIC. This distinct input pattern, combined with its long-latency sound responses, clearly distinguishes cMGv from rMGv. Thus, while our results confirm the cMGv→A2 projection^34^, we propose that both cMGv and its target A2 should be considered components of the secondary non-lemniscal pathway. Collectively, our findings and previous studies establish multiple parallel pathways within both the lemniscal and non-lemniscal divisions of the auditory system.

Secondary sensory cortices are generally thought to receive inputs from primary cortices through both cortico-cortical and cortico-thalamo-cortical pathways^23,24^. Consistent with this organization, A2 receives inputs from A1 both directly^59,60^ and indirectly via MGd^34,61,62^ (Fig. 3e). Altogether, our results indicate that A2 serves as a convergence site for at least three thalamocortical pathways: the non-lemniscal MGm/BIC input to L6, the non-lemniscal cMGv input to L4, and the canonical MGd input to L4 that indirectly conveys lemniscal signals from A1. In our retrograde tracing experiments from A2 L4, we found nearly five times more input cells in cMGv than in MGd. Nevertheless, since our electrophysiological recordings in A2 L4 reflect integrated signals from both thalamic regions with slow kinetics, further studies will be crucial to determine how these two L4 inputs differentially shape A2 sound responses. Understanding how these three thalamocortical pathways contribute to the specialized sound response properties of A2^41,42,63–65^ remains an important direction for future research.

The existence of parallel ascending pathways that directly reach higher-order sensory cortices, bypassing primary areas, appears to be a common motif across sensory systems. In the visual system, the higher-order postrhinal cortex (POR) receives visual information in L4 via a tecto-pulvinar pathway, parallel to the canonical geniculo-striatal input to the primary visual cortex (V1)^1,4,7^. Similarly, in the somatosensory system, while the principal trigeminal nucleus (Pr5) signals to the primary somatosensory cortex (S1) via the ventral posterior medial nucleus (VPM), the spinal trigeminal nucleus (Sp5i) transmits information to L4 of the secondary somatosensory cortex (S2) through the rostral posterior medial thalamus (POm)^3^. Our discovery of an ascending auditory pathway to A2 L4 through cMGv, which we have identified as part of the non-lemniscal thalamus, thus indicates a conserved organizational principle across sensory modalities: alternative pathways transmit sensory information to L4 of higher-order cortices through non-primary thalamic nuclei, complementing the primary pathways. This accumulating evidence challenges the classical hierarchical model of sensory processing and suggests that cortical circuits implement both hierarchical and parallel processing schemes simultaneously.

### Cortical layer-selective inputs from non-lemniscal MGN neurons

Neurons in secondary thalamic nuclei have traditionally been considered to project broadly across cortical layers^35^. Contrary to this assumption, our observation of non-lemniscal short-latency inputs specifically in L6, but not in L2/3–L5, indicates that short-latency MGm/BIC neurons actually exhibit cortical layer-selective projections. We do not rule out the presence of additional projections to L1, as linear probe recordings are not sensitive to L1 neuronal firing, and input to L1 local interneurons, due to their dendritic geometry, may not generate significant extracellular fields and detectable CSD sinks. Nevertheless, the layer selectivity is particularly evident in A2, where the L6 input precedes responses in L2/3, L4, and L5a by approximately 5 ms—far greater than would be expected from axonal conduction delay alone. This finding also rules out the possibility that the L6 short-latency input originates from collaterals of L4-projecting thalamocortical axons.

Our results suggest that what previously appeared to be layer-nonspecific projections may have resulted from bulk labeling of multiple, functionally distinct subpopulations, each with its own specific layer targets. Supporting this idea, small-volume anterograde tracer injections into MGm and SG have revealed projection patterns favoring L6 and L1, or L6 alone^66^—patterns clearly distinct from the L1-and L4-preferring projections of calretinin-expressing MGd neurons^17,67^. Future identification of genetic markers that selectively label these functional MGN subpopulations will be crucial for better characterizing the parallel non-lemniscal pathways that deliver information to individual cortical layers.

### Short-latency responses in the non-lemniscal auditory thalamus

We found that the short-latency sound responses observed in L6 of both A1 and A2 originate from the MGm and BIC, which themselves exhibit short-latency responses. The MGm is characterized by its heterogeneous cytoarchitecture^68^ and electrophysiological properties^8,46,47^, earning the description as “the most enigmatic of the auditory thalamic nuclei^69^”. Response latencies in this region are also heterogeneous, and a subpopulation with latencies comparable to those of primary MGv neurons has been observed across species^8,45–47^. Our identification of short-latency responses specifically in neurons with short P-T interval spikes further supports the presence of functionally distinct subpopulations within MGm. In contrast, neurons with long P-T intervals showed significantly longer-latency sound responses similar to MGd neurons, suggesting that they may be driven preferentially by DCIC or cortical feedback, potentially serving as a conduit between primary and secondary auditory cortices. Determining whether these two MGm subpopulations show distinct connectivity and functional roles in sound processing will be an interesting topic of future research.

Direct projections from the cochlear nucleus to MGm, documented in previous studies^48,49^, could potentially explain the short-latency sound responses observed in this thalamic division. In line with these studies, our disynaptic rabies tracing from cortical L6 revealed sparse projections from the cochlear nucleus to starter cells in MGm and BIC. However, the fraction of cochlear nucleus input cells was only 0.3% of those originating in the inferior colliculus. While we cannot rule out the possibility that this sparse direct projection contributes to MGm responses, our observation of short-latency responses in the rostral ECIC suggests a predominant contribution from a more conventional cochlear nucleus→inferior colliculus→MGm pathway.

In addition to MGm, we unexpectedly identified the rostral portion of BIC as another relay station projecting to L6 of both A1 and A2. BIC is an understudied region that extends along the rostrocaudal axis between the inferior colliculus and the MGN. Traditionally, it has been considered an extension of the inferior colliculus, as the name suggests, due to its anatomical connections with the inferior colliculus and the superior colliculus^70–72^. However, our findings lead us to propose that the rostral BIC more closely aligns with the MGN for three reasons: First, the rostral BIC contributes direct projections to the auditory cortex (Fig. 3). Second, tracing the projections of the thalamic reticular nucleus (TRN), an inhibitory nucleus that innervates the thalamus^73^, revealed its axons innervating the rostral, but not caudal BIC (Supplementary Fig. 3a,b). This implicates the rostral BIC as part of the canonical thalamic circuit that includes the TRN. Third, calbindin-1, a marker that strongly labels non-lemniscal MGN, is also expressed in the rostral BIC, distinguishing it from the caudal BIC (Supplementary Fig. 3c)^67,74^. Collectively, these observations suggest that BIC should be subdivided into rostral and caudal subdivisions, with the rostral region having a close functional association with MGN.

### Short- and long-latency ascending pathways in the non-lemniscal inferior colliculus

Our retrograde tracing from cMGv and MGm^L6^ revealed that cMGv receives inputs predominantly from more medial and caudal domains of the inferior colliculus shell (primarily DCIC), while MGm^L6^ is predominantly innervated by more rostral domains (primarily ECIC). These findings not only support the non-lemniscal identity of cMGv, but also reveal two overlapping but distinct non-lemniscal sources for the slow and fast sound responses transmitted to A2 L4 and L6, respectively. Our electrophysiological recordings across the inferior colliculus further supported these findings, demonstrating corresponding response latencies in these non-lemniscal subdivisions: slow responses in the caudomedial DCIC and fast responses in the rostrolateral ECIC.

These observations align with recent findings in guinea pigs, which also reported short-latency responses in the rostral ECIC^75^, suggesting the conservation of this structural organization across species. Both the guinea pig study and our mouse data demonstrated caudomedial-to-rostrolateral gradients of response latencies within CIC and ECIC without significant differences between rostral CIC and rostral ECIC. The short-latency responses in rostral ECIC may have gone unnoticed in earlier research because previous characterizations of the inferior colliculus focused primarily on its caudal half. Our findings thus challenge the traditional dichotomy of fast lemniscal and slow non-lemniscal pathways and instead support a revised circuit architecture featuring parallel fast pathways ascending through rostral domains of both lemniscal and non-lemniscal inferior colliculus.

### Roles of short-latency L6 inputs in cortical processing

Short-latency inputs to L6 likely serve multifaceted roles in cortical sensory processing. These direct ascending pathways, which bypass superficial layers, may rapidly activate the output layers of the cortical circuit, allowing the animal to react promptly to incoming stimuli. Additionally, given that L6 activity can have both excitatory^76^ and inhibitory^77,78^ influences on more superficial layers, these deep-layer inputs could modulate sensory information processing throughout the cortical column. While our analysis indicates that all L6 functional cell types receive these ascending inputs (but see^79^), further investigation of input strengths across individual cell types may provide deeper insights into how L6 inputs shape cortical information processing.

We found that non-lemniscal ascending inputs to L6 in both A1 and A2 possess broader frequency tuning compared to the lemniscal input to A1 L4. This less selective, more integrative nature of non-lemniscal L6 input may allow the transmission of spectrotemporally complex auditory information to the cortex. Alternatively, rather than conveying detailed spectral information, these short-latency signals may instead serve as an early alerting mechanism, informing cortical circuits of incoming stimuli and priming them for subsequent, more detailed sensory processing. In either scenario, the distinct tuning properties highlight the functional differentiation between lemniscal A1 L4 input and non-lemniscal L6 inputs.

Notably, non-lemniscal subcortical structures receive extensive top-down projections^3,13,15,16^, positioning these pathways to dynamically modulate cortical sensory processing based on behavioral contexts. Through selective modulation of these ascending signals, the brain may flexibly adjust how incoming sensory information is interpreted and prioritized. Future research should investigate the specific information carried by these parallel pathways and determine how top-down influences shape these ascending inputs. Ultimately, understanding the interplay between direct L6 inputs, top-down modulation, and the broader cortical activities triggered by lemniscal inputs will be crucial for unraveling the full complexity of cortical sensory processing.

## Methods

### Animals

Mice were at least 6 weeks old at the time of experiments. The strains used include: C57BL/6J (JAX000664), B6N-Cdh23^tm2.1Kjn^/Kjn (JAX 018399), Sst^tm2.1(cre)Zjh^/J (Sst-Cre; JAX 013044), Pvalb^tm1(cre)Arbr^/J (PV-Cre; JAX 017320), and Tg(Rbp4-Cre)KL100Gsat/Mmucd (Rbp4-Cre; MMRRC 037128-UCD). Both female and male mice were used. The animals were housed at 21°C and 40% humidity, with ad libitum access to food pellets and water. They were kept under a reverse light cycle (12– 12 h), and all experiments were conducted during their dark cycle. All experimental procedures were approved and conducted in accordance with the Institutional Animal Care and Use Committee at the University of North Carolina at Chapel Hill and the guidelines of the National Institutes of Health.

### Auditory stimuli

Auditory stimuli were calculated in Matlab (MathWorks) at a 192 kHz sample rate and delivered via a free-field electrostatic speaker (ES1; Tucker-Davis Technologies). The speakers were calibrated across a range of 2–64 kHz, and stimulus voltages were adjusted to give a flat response (±1 dB) across this range. Stimuli were delivered to the ear contralateral to the imaging or recording site. Stimulus delivery was controlled by Bpod (Sanworks), running in Matlab.

For areal mapping with intrinsic signal imaging, 3, 10, and 30 kHz pure tones (75 dB SPL, 1-second duration) were presented at 30-second intervals for 5–20 trials. For electrophysiological recordings, mice were presented with either 100 ms pure tones covering 17 frequencies (4–64 kHz, log-spaced) at 25, 40, 55, and 70 dB SPL or 200 ms pure tones covering 9 frequencies (4–64 kHz, log-spaced) at 70 dB SPL. All pure tones had a 5-ms linear rise and fall at their onset and offset. Tones were presented in random order at 0.7–2-second intervals, with each stimulus repeated across nine trials. Both 9- and 17-frequency datasets were used to determine tone response latencies, while only 17-frequency datasets were used to assess frequency tuning broadness. Click stimuli were generated as 0.1-ms monopolar rectangular pulses and presented across 200 trials at 0.5-second intervals.

### Intrinsic signal imaging

Intrinsic signal imaging was conducted to locate the auditory cortical areas, following a previously described protocol^39^. Signals were acquired using a custom tandem lens macroscope comprising Nikkor 35 mm 1:1.4 and 135 mm 1:2.8 lenses, attached to a 12-bit CCD camera (DS-1A-01M30, Dalsa), placed in a sound-attenuating enclosure (Gretch-Ken Industries). Mice were anesthetized with isoflurane (0.8– 2%) vaporized in oxygen (1 L/min) and maintained on a heating pad at 36°C. The muscle overlying the right auditory cortex was removed, and a custom stainless steel head bar was secured to the skull with dental cement. The brain surface was imaged through the skull, which was kept transparent with phosphate-buffered saline (PBS). Mice received a subcutaneous injection of chlorprothixene (1.5 mg/kg) prior to imaging. Images of surface vasculature were captured under green LED illumination (530 nm), and intrinsic signals were recorded at 16 Hz using red illumination (625 nm). Images of reflectance were acquired at a resolution of 717 × 717 pixels (2.3 × 2.3 mm). Images collected during the response period (0.5–2 seconds from sound onset) were averaged across trials and normalized to the average image during the baseline period. The images were Gaussian-filtered and thresholded for visualization. Individual auditory areas, including A1, AAF, VAF, and A2, were identified based on their characteristic tonotopic _patterns_^39,58,65,80^.

### Electrophysiology

For A1 and A2 recordings, mice were first implanted with a head bar, and auditory cortical areas were mapped using intrinsic signal imaging 1-3 days before recording. On the day of recording, a small craniotomy (<0.3 mm) and durotomy were performed at the mapped location of A1 or A2. Awake mice were head-fixed, and either a 64-channel silicon probe (ASSY-77-H3, sharpened, Cambridge Neurotech) or a manually sharpened 384-channel Neuropixels 1.0 probe (imec) was slowly inserted perpendicular to the brain surface at approximately 1 µm per second. Probes were painted with DiI or DiO (in ethanol) and were allowed to dry prior to use. Spikes were monitored during probe insertion, and the probe was advanced until its tip reached the white matter, indicated by the absence of spikes. A reference electrode was placed on the dura above the visual cortex. After insertion, the probe was allowed to settle for at least one hour before data collection. During recordings, mice sat quietly, with occasional bouts of whisking and grooming, in a loosely fitted plastic tube lined with fleece fabric for comfort and attenuation of scratching noise. The stage was placed within a sound-attenuating enclosure (Gretch-Ken Industries or custom-built). Extracellular voltage signals were amplified, digitized, and acquired at 20 kHz (64-channel probes) or 30 kHz (Neuropixels) using the OpenEphys system (https://open-ephys.org). Post hoc identification of probe trajectories was achieved by examining DiI or DiO fluorescence in brain sections counterstained with DAPI.

In some A1 experiments, after cortical recordings were completed, the probe was advanced deeper to reach MGN. In other MGN recording experiments, mice received head bar implantation without intrinsic signal imaging, and MGN was stereotaxically targeted at coordinates around (A: −3.16 mm, L: 1.75 mm, V: 3.5 mm) at 24 degrees from vertical. All coordinates are given relative to bregma (A: anterior; L: lateral; V: ventral).

For caudal inferior colliculus recordings, mice were implanted with a head bar without intrinsic signal imaging. The skull was thinned until the inferior colliculus was visible. On the recording day, a small craniotomy and durotomy were performed, and the probe was visually guided to the target division of the inferior colliculus. Rostral inferior colliculus was stereotaxically targeted at coordinates around (A: −4.7 mm, L: 1.5 mm, V: 2.5 mm) at 24 degrees from vertical.

### Analysis of electrophysiology data

Single- and multi-unit activity was isolated using Kilosort 2.5 software^81^ and the spike-sorting graphical user interface Phy^82^. The initial automated classification was performed using the Bombcell spike classification toolbox^83^, followed by the inspection and editing of individual clusters to remove artifacts and duplications. Multi-unit spikes were calculated by aggregating all spikes within each channel (Figures 1n,o, 4c-e, 6c-g, Supplementary Figures 1a,b, 4), layer (Figures 1g,i,k,m, 2, Supplementary Fig. 2), or region (Figures 4b and 6b). For spike waveform analyses (Fig. 4f-i, Supplementary Figures 1c-e, 5), individual spike clusters sorted by Kilosort were used after noise removal. Fast-spiking units were classified based on their small P-T interval (≤ 0.4 ms). For CSD analysis, LFP at individual channels was low-pass filtered at 1000 Hz, averaged across sound presentation trials, and processed using inverse CSD calculation^84^. In Neuropixels recordings, CSD signals were computed separately for the four columns of channels and then averaged. CSD visualization involved normalization to peak sink magnitudes across channels and Gaussian smoothing along the cortical depth (σ = 0.02×cortical thickness). Peri-stimulus time histograms (PSTHs) were constructed using 0.5-ms bins and summed across sound presentation trials. PSTH visualization involved normalization to the PSTH peak across channels and Gaussian smoothing along the cortical depth (σ = 0.02×cortical thickness) and time (σ = 1 ms).

In cortical recordings, layer boundaries were determined using a combination of CSD signals, spikes, and LFP coherence across channels^40^. The cortical surface (pia) was identified by a sharp reduction in LFP amplitude for shallower sites, and the white matter was marked by the absence of spikes. In CSD signals, L4 channels (L4_CSD_) were identified as the early, strong sink in the middle portion of the cortical column. LFP coherence was calculated using Pearson correlation coefficients between channel pairs from pia to white matter during spontaneous activity (10 minutes) or pure tone exposures (8 minutes). Full-bandwidth LFP (1–64 Hz) typically revealed four clusters of coherent channels, corresponding to L1, L2-4, L5, and L6. In some cases, delta-band (1–4 Hz) or beta-gamma band (32–64 Hz) LFP provided clearer cluster separation. Application of k-means clustering to LFP traces of individual channels (k = 4; seeds at supragranular center, L4 center, and upper/lower halves of subgranular layer, estimated from CSD) gave consistent results with the LFP coherence and was used to define channels for L1_LFP_, L2-4_LFP_, L5_LFP_, and L6_LFP_. The final layer boundaries were determined as follows: L1-L2 and L5-L6 borders were set between L1_LFP_/L2-4_LFP_ and L5_LFP_/L6_LFP_ borders, respectively. The L2/3-L4 border was defined by the top channel of L4_CSD_. The L4-L5 border was determined as the mean position between L4_CSD_ bottom and L2-4_LFP_ bottom channels. Average positions of layer borders across all mice (normalized to 0 = pia; 1 = white matter top) were: A1: L1-L2 = 0.182, L2/3-L4 = 0.293, L4-L5 = 0.485, L5-L6 = 0.758; A2: L1-L2 = 0.182, L2/3-L4 = 0.263, L4-L5 = 0.475, L5-L6 = 0.768. For visualization of averaged CSD and PSTH across mice, individual heat maps were normalized to peak magnitudes and morphed to these averaged layer boundaries.

First-spike latencies of individual multi-units (Figures 1n,o, 4c-e, 6c-g, Supplementary Figures 1a,b, 4) were analyzed only for significantly sound-responsive units. Unit-sound pairs were judged as significantly responsive if they fulfilled two criteria: 1) PSTH had to exceed a fixed threshold at the same time bin for more than one-third of trials. 2) Trial-averaged PSTH had to exceed a fixed threshold value. The threshold for excitation (3.7 × standard deviation of the baseline period) was determined using ROC analysis to achieve 90% true positive rate in tone-evoked responses. First-spike latencies were determined as the first significant deviation of spike rate from spontaneous rate, following established methods^85^. PSTHs were constructed using 1-ms bins, with a threshold probability of *p* = 1.0×10^−3^ for detecting deviations. For layer- and region-based analyses (Figures 1f-m, 4b, 6b), response latencies were defined simply as the time when population CSD or PSTH signals, averaged across channels, exceeded the threshold of 3 × standard deviation of the baseline period. Tone response latencies were calculated using trials with the best frequency (BF) and its two adjacent frequencies. BF for each recording site was determined as the tone frequency that triggered the maximum spike response, calculated as the sum of spikes across all cortical layers during the first 50 ms after tone onset.

To quantify layer-specific inputs while minimizing across-layer propagation, L6-preference index (Fig. 1p,q) and tuning broadness (Fig. 2 and Supplementary Fig. 2) were calculated using brief time windows around response onsets. The L6-preference index was calculated as (L6 response – L4 response) / (L6 response + L4 response) and ranged from -1 to 1. Response amplitudes were quantified as the mean CSD sink or PSTH above baseline during a 5-ms window following the layer-specific response latency, which was determined for each mouse as described above. For frequency tuning broadness analysis, fixed quantification windows were used across all frequencies since response latencies to non-preferred tones were often indeterminate. The windows were 5-15 ms after tone onset for A1 L4, A1 L6, and A2 L6, and 10-20 ms for A2 L4. Response amplitudes to 70 dB SPL tones were fitted with Gaussian curves, from which full-width half-maximum (FWHM) values were derived. Only fits with R-square values larger than 0.4 were included in the analysis. Tuning curves were centered at the BF of the recording site and normalized to the maximum response amplitudes for each layer and mouse.

In MGN recordings, division boundaries were determined by mapping DiI- or DiO-marked probe trajectories to the Paxinos Brain Atlas^44^ using a custom Matlab code. Recording coordinates were established by comparing coronal brain sections with the Brain Atlas to determine rostrocaudal position, thalamic entry point coordinates (dorsoventral and mediolateral), and probe angle. The channel at the thalamic entry point was identified based on the appearance of spikes. Coordinates for deeper channels were then calculated from the thalamic entry point according to the probe angle and an inter-channel spacing of 20 µm. In some experiments, MGN division borders were further verified using mice expressing fluorophores in TRN axons (PV-Cre and Sst-Cre mice) or A1 corticothalamic projections (Rbp4-Cre mice). The boundary between the rostral and caudal subdivisions of MGv was set at −3.30 mm from bregma. However, we note that the observed latency differences between rMGv and cMGv (Fig. 4e) are likely underestimated due to the complex three-dimensional nature of this boundary (Fig. 3f)^34^. Due to the limited number of SG units and unclear borders between structures, we combined SG with MGm for quantification, except in Supplementary Fig. 4. We excluded the marginal zone of the medial geniculate (MZMG) from MGv, as it shows distinctly different activity patterns (Supplementary Fig. 4) and gene expression^34^. For visualization in Fig. 4c, 4e, and Supplementary Fig. 4a-c, small jitter was added to the rostrocaudal coordinates to minimize overlaps.

For inferior colliculus recordings, division boundaries were determined by mapping probe trajectories onto the Allen Common Coordinate Framework 2017 using Matlab codes modified from a publicly available repository (https://github.com/petersaj/AP_histology). The boundary between rostral and caudal subdivisions of the inferior colliculus was set at −4.9 mm from bregma, corresponding to the midpoint of the structure’s rostrocaudal extent. Note that our definition of rostral ECIC was more conservative than the Paxinos Brain Atlas, which defines all inferior colliculus rostral to −4.9 mm as ECIC^44^. Using the Paxinos Brain Atlas would have classified our rostral CIC as part of rostral ECIC, resulting in even shorter latencies in rostral ECIC.

We performed k-means clustering on multi-unit spike waveforms sorted by Kilosort 2.5 and manually cleaned in Phy2. For each multi-unit, we computed a median spike waveform across trials, and the waveform at the channel with the peak amplitude was used for clustering. This waveform was smoothed with a 5-sample moving mean and normalized by Z-scoring. For each multi-unit, a waveform vector was constructed from 82 samples surrounding the waveform trough (40 samples before and 41 samples after the trough). We concatenated all such vectors (number of multiunits × 82) and performed k-means clustering with 10 repetitions. Elbow point analysis indicated optimal cluster numbers of 2 for MGm (Fig. 4f-i) and BIC (Supplementary Fig. 5b-d) and 3 for cortical L6 (Supplementary Fig. 1c-e). Because Cambridge Neurotech and Neuropixels probes have different sampling rates, cortical clustering was performed exclusively on data from Cambridge Neurotech recordings (23 mice in A1, 14 mice in A2). All MGN recordings were conducted using Cambridge Neurotech probes.

### Anatomical tracing

Cortical area- and layer-specific retrograde tracing was performed following transcranial mapping of auditory cortical areas using intrinsic signal imaging. Retrograde tracers (fluorophore-conjugated cholera toxin B subunit: CTB-488, 555, or 594) were targeted to either A1 or A2 at various frequency domains. While some studies consider VAF to be a ventral branch of A1, we adopted a conservative approach and focused on a narrowly defined A1, excluding the mid-high frequency domains of VAF. Through small craniotomies, 0.5% CTB solution (40 nL at 10 nL/min) was delivered using beveled glass pipettes (20 µm external diameter) at depths of 320–360 µm for L4 and 700–800 µm for L6 (A2 and A1, respectively). Four to seven days after injection, mice were transcardially perfused with PBS, followed by 4% paraformaldehyde in PBS under isoflurane anesthesia. Brains were extracted, fixed overnight in 4% paraformaldehyde in PBS, cryoprotected in 30% sucrose in PBS, and coronally sectioned at 40 μm thickness using a freezing microtome. Sections were mounted on slide glasses, counterstained with DAPI, and imaged using either a confocal microscope (Olympus FV3000RS) or an optical dissection fluorescence microscope (Zeiss Axio Observer 7). MGN division boundaries were identified using DAPI staining and tracer-labeled neuropil signals and corroborated by comparing brain sections with calbindin and SMI-32 expression patterns (https://connectivity.brain-map.org/static/referencedata)^34^. The rBIC boundaries with MGm and caudal BIC were set at −3.5 mm and −4.1 mm from bregma, respectively. For each area-layer pair, the fraction of input cells in each MGN division (Fig. 3d,e) was calculated relative to the total MGN input cells in each mouse. The rostrocaudal distributions (Fig. 3f,g) were calculated as fractions relative to the total input cells within each specific pathway, and the averaged data across mice were displayed.

Rabies disynaptic retrograde tracing from L6 (TRIO method)^86^ was conducted following transcranial mapping of auditory cortical areas with intrinsic signal imaging. Through a small craniotomy, AAVretro-syn-Cre (Addgene; 30 nL at 10 nL/min) was injected into L6 at depths of 700 µm (A2) or 800 µm (A1). During the same surgery, a mixture of AAV9-CAG-Flex-mKate-T2A-TVA and AAV9-CAG-Flex-mKate-T2A-CVS-N2cG (UNC BRAIN Initiative Viral Vector Core; 300 nL at 20 nL/min) was stereotaxically targeted to the MGN at coordinates (A: −3.4 mm, L: 1.9 mm, V: 3.4 mm). After fourteen days, EnvA-rabies-CVS-N2c(ΔG)-H2B-EGFP (UNC BRAIN Initiative Viral Vector Core; 400 nL at 20 nL/min) was injected at the same MGN location. Seven days after the second injection, tissue preparation and imaging were performed as described above. One mouse (the data point with the lowest cell count in Supplementary Fig. 7b) received AAV8-hSyn-Flex-TVA-P2A-GFP-2A-oG and EnvA-ΔG-rabies-mCherry (Salk Institute Viral Vector Core) with a four-day wait period following the rabies injection. Control animals received MGN viral injections without the cortical AAVretro-syn-Cre injection. Starter cells (EGFP^+^/mKate^+^) and input cells (EGFP^+^) were counted semi-automatically using QuPath software (https://qupath.github.io/). Cells were manually assigned to inferior colliculus divisions by comparing fluorescence tracer signals with the Allen Common Coordinate Framework v3^87^ and verified against Nos1 expression patterns (https://connectivity.brain-map.org/static/referencedata).

For monosynaptic retrograde tracing from MGN subdivisions, a large craniotomy (approximately 3– 4 mm in diameter) was made over the right auditory cortex. Following durotomy, the cortex and hippocampus above the MGN were aspirated to expose the MGN and LGN surfaces. Using sharpened glass pipettes, a retrograde tracer (0.5% CTB-488, 594, or red Retrobeads IX, Lumafluor; 30 nL, 10 nL/min) was injected into MGd (200 µm deep), cMGv (250 µm deep), or rMGv (500 µm deep through the LGN). After injection, the brain cavity was filled with gel foam, and the skull flap was placed back to cover the craniotomy. Four to seven days after injection, tissue preparation and imaging were performed as described above. The fraction of input cells in each inferior colliculus division (Fig. 5c) was calculated relative to the total inferior colliculus input cells in each mouse. The rostrocaudal distributions (Fig. 5g) were calculated as fractions relative to the total input cells within each specific pathway, and the averaged data across mice were presented.

For monosynaptic retrograde tracing from the CIC, DCIC, and caudal ECIC, we thinned the skull over the inferior colliculus to visualize the structure. Following a small craniotomy above the target division, we injected a retrograde tracer (0.5% CTB-488, 594, 0.5–1% fluorophore-conjugated wheat germ agglutinin WGA-488, 594, or 647; 30–40 nL at 10–20 nL/min). The depth of injection was 600 µm for DCIC, 1000–1200 µm for CIC (vertical), and 600–900 µm for ECIC (24° from vertical). Rostral ECIC was stereotaxically targeted at the coordinates (A: −4.6 mm, L: 1.5 mm, V: 2.35 mm). Three to five days after injection, tissue preparation and imaging were performed as described above. Input cells were manually assigned to cochlear nucleus divisions using DAPI staining as a guide. Within the ventral cochlear nucleus, we defined AVCN and PVCN as regions rostral and caudal to the nerve root, respectively.

TRN axon projections to MGN were visualized by injecting AAV8-nEF-Con/Foff2.0-ChRmine-oScarlet into the auditory TRN (A: −1.85 mm, L: 2.30 mm, D: 3.50 mm) of Sst-Cre mice. Sst-expressing neurons in TRN project their axons preferentially to the higher-order thalamus^73^.

### Statistics

Sample sizes were not predetermined using statistical methods but were selected based on standards commonly used in the field. All “*n*” values refer to the number of mice or units, as identified in the text. All key results were replicated across multiple mice. Mice were randomly assigned to experimental groups. While the experiments were not performed blind, experimenters were blinded to trial types and timings during spike sorting in electrophysiology. Data are presented as individual data points along with median or mean ± SEM, as specified in figure legends. Statistical differences between conditions were evaluated using a two-sided Wilcoxon rank-sum test. P-values were corrected for multiple comparisons using the Bonferroni correction. Correlation coefficients were calculated as Pearson’s correlation using Matlab’s corrcoef function. Two-way ANOVA was used to evaluate the effects of independent factors.

## Data availability

All data generated in this study will be deposited in the DANDI Archive. Source data for all figures will be provided in this paper as a supplementary data file attached to this paper.

## Code availability

Custom Matlab codes used in this study will be made available from the corresponding author upon request.

## Acknowledgements

We thank Kendall Hutson and the members of the Kato Lab for their advice throughout the project and comments on the manuscript. This work was supported by National Institute on Deafness and Other Communication Disorders (R01DC017516: H.K.K.), National Institute of Neurological Disorders and Stroke and BRAIN Initiative (RF1NS128873: H.K.K. and P.B.M.; F31-NS111849: A.M.K.), and Japan Society for the Promotion of Science (K.O.). Part of the microscopy was performed at the UNC Neuroscience Microscopy Core (RRID: SCR_019060), supported, in part, by funding from the NIH-NINDS Neuroscience Center Support Grant P30 NS045892 and the NIH-NICHD Intellectual and Developmental Disabilities Research Center Support Grant P50 HD103573.

## Author contributions

M.M.G., H.T., P.B.M., and H.K.K. designed the project. M.M.G., P.R.D., H.C.A., and H.K.K. performed tracing experiments. M.M.G., A.M.K., K.O., and H.K.K. performed electrophysiology experiments. M.M.G., A.M.K., K.O., H.T., M.K., P.B.M., and H.K.K. performed data analysis. M.M.G. and H.K.K. wrote the manuscript with inputs from all the authors.

## Additional information

Correspondence and requests for materials should be addressed to Hiroyuki K. Kato.

