## Supplementary Information for "Noncanonical Short-Latency Auditory Pathway Directly Activates Deep Cortical Layers"

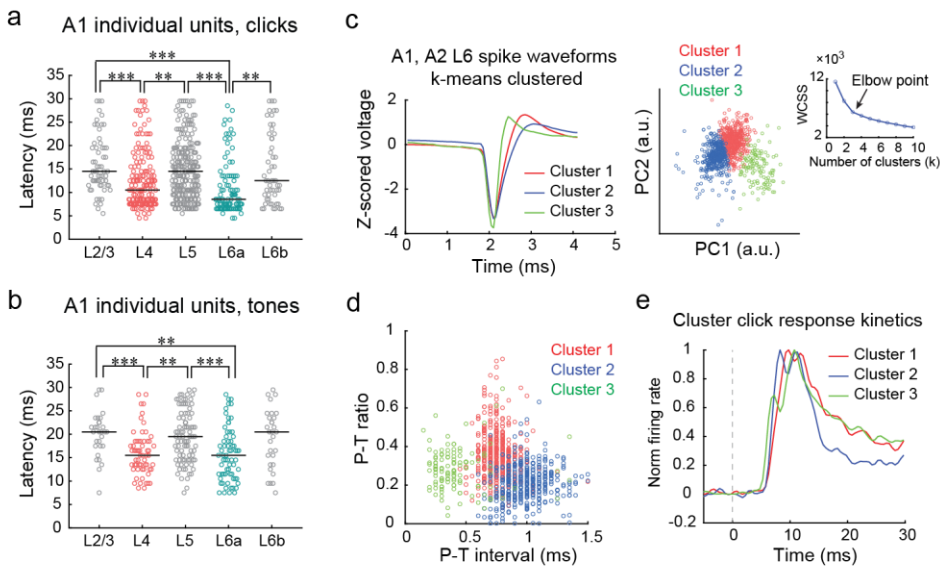


**Supplementary Figure 1. Additional data characterizing sound-evoked spike responses in A1 and A2.** **(a)** First-spike latencies of A1 click responses in individual units by layer ($n$ = 31 mice; **$p$ < 0.01, ***$p$ < 0.001, two-sided Wilcoxon rank sum test with Bonferroni correction). **(b)** First-spike latencies of A1 BF tone responses in individual units by layer. **(c)** Left, averaged spike waveforms for units in Cluster 1 (red), Cluster 2 (blue), and Cluster 3 (green). Middle: three clusters of L6 spikes in PC space. Right: elbow point analysis (optimal cluster number = 3). WCSS: within-cluster sum of squares. A1: $n$ = 31 mice, A2: $n$ = 16 mice. **(d)** Distributions of P-T interval and P-T ratio for individual L6 clusters. **(e)** Click-triggered PSTHs for individual L6 clusters. All three clusters show short-latency responses.

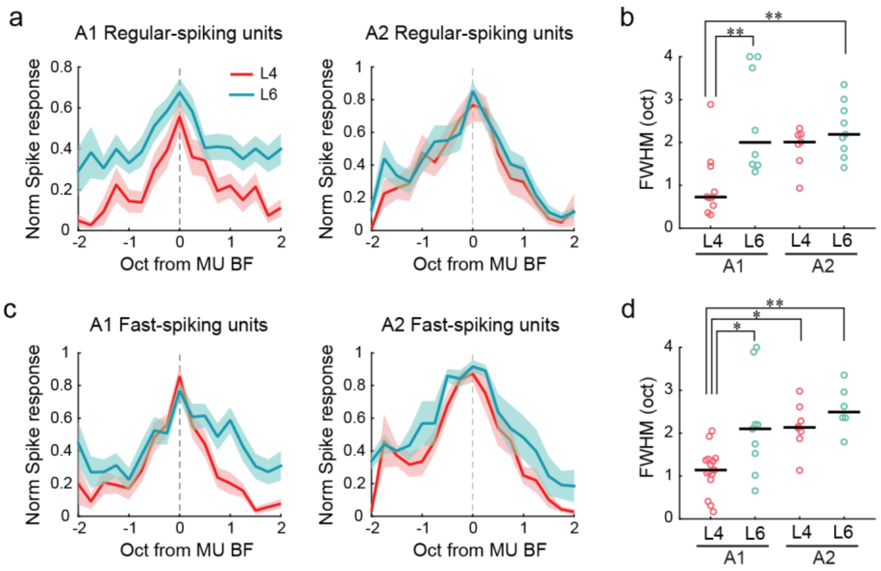


**Supplementary Figure 2. Both regular-spiking and fast-spiking units in L6 show broad frequency tuning. (a)** Normalized frequency tuning curves of regular-spiking units in L4 and L6 of A1 (left) and A2 (right), centered at the BF of the recording site (A1: $n$ = 20 mice; A2: $n$ = 9 mice). Lines: mean. Shading: SEM. **(b)** Summary of full-width half-maximum (FWHM) of tuning curves for regular-spiking units (*$p < 0.05$, **$p < 0.01$, two-way ANOVA followed by Tukey's HSD test). Black bars: median. **(c-d)** Same as (a-b), but for fast-spiking units.

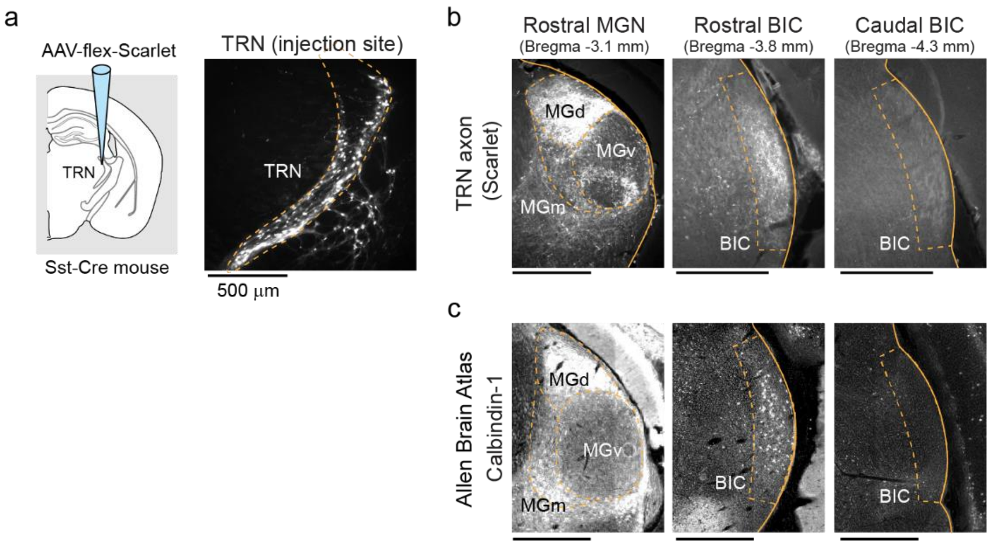


**Supplementary Figure 3. Rostral BIC exhibits characteristics of the non-lemniscal MGN. (a)** Left: schematic of anterograde tracer injection into the auditory TRN. Right: coronal section showing Scarlet-expressing neurons in the auditory TRN. **(b)** Representative coronal sections showing TRN axon terminals at rostral MGN, rostral BIC, and caudal BIC levels. TRN axons are present in rostral but not caudal BIC. Similar results obtained in four mice. **(c)** Calbindin-1 expression patterns at rostral MGN, rostral BIC, and caudal BIC levels. Data from the Allen Mouse Brain Atlas. https://connectivity.brain-map.org/static/referencedata/experiment/siv/100142355?imageId=102167376&imageType=CALB1. Rostral MGN: image 139; Rostral BIC: image 154; Caudal BIC: image 162. Calbindin-1 is expressed in rostral but not caudal BIC.

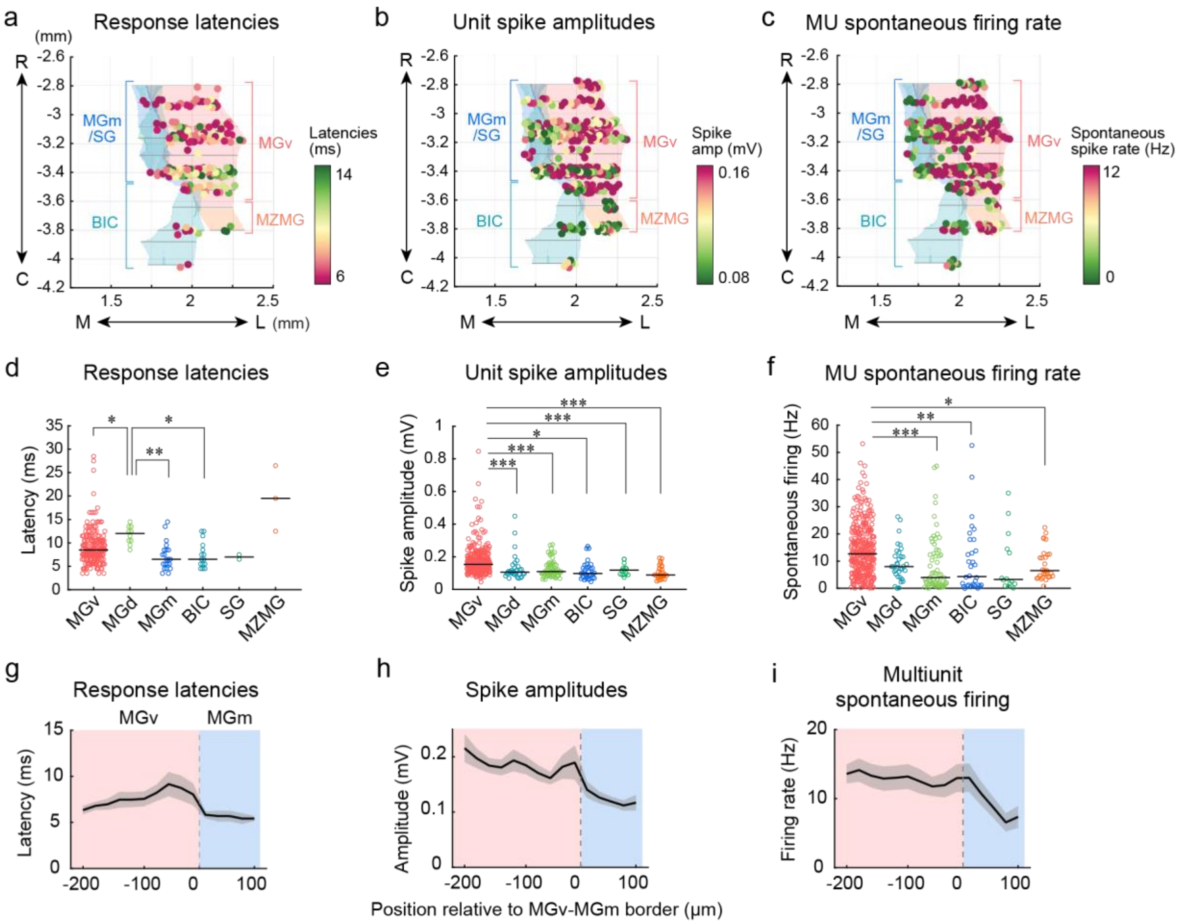


**Supplementary Figure 4. Additional data characterizing multiunit activity in MGN divisions. (a)** Click response latencies at reconstructed recording sites across MGN divisions. Dorsal view of 3-D reconstructions from 26 mice. MGd excluded to visualize the transition between cMGv and MGm/BIC. Data same as Fig. 4c. **(b)** Spike amplitudes at reconstructed recording sites across MGN divisions. MGv shows larger spike amplitudes than non-lemniscal divisions. **(c)** Multiunit spontaneous firing rates at reconstructed recording sites across MGN divisions. Channels in MGv show higher rates than non-lemniscal divisions. **(d–f)** First-spike latencies of click responses **(d)**, spike amplitudes **(e)**, and spontaneous firing rates **(f)** in individual units for each MGN division (*$p < 0.05$, **$p < 0.01$, ***$p < 0.001$; two-sided Wilcoxon rank sum test with Bonferroni correction). The data for MGv, MGd, MGm, and BIC in (d) are the same as Fig. 4d. Black bars: median. **(g–i)** Changes in first-spike latencies of click responses **(g)**, spike amplitudes **(h)**, and spontaneous firing rates **(i)** around the MGv-MGm transition border ($n$ = 13 mice with recording track crossing the MGv-MGm border). Red and blue areas: MGv and MGm channels, respectively. Solid line: mean. Shading: SEM.

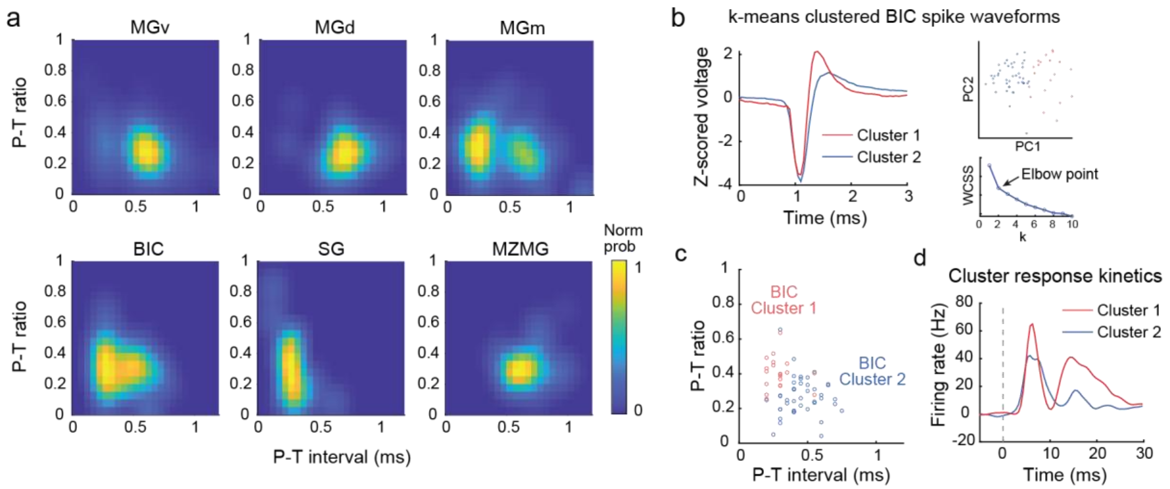


**Supplementary Figure 5. Additional data for neural subpopulation analyses in MGN. (a)** Two-dimensional heat maps showing the probability distribution of P-T intervals (x-axis) and P-T ratio (y-axis) for each MGN division. Data for MGv, MGd, MGm, and BIC are the same as Fig. 4f. (b-d) k-means clustering of BIC spike waveforms. **(b)** Left: averaged spike waveforms for units in Cluster 1 (red) and Cluster 2 (blue). Top right: two clusters of BIC spikes in PC space. Bottom right: elbow point analysis (optimal cluster number = 2). WCSS: within-cluster sum of squares. **(c)** Distributions of P-T interval and P-T ratio for individual BIC clusters. **(d)** Click-triggered PSTHs for individual BIC clusters. Both clusters show short-latency responses.

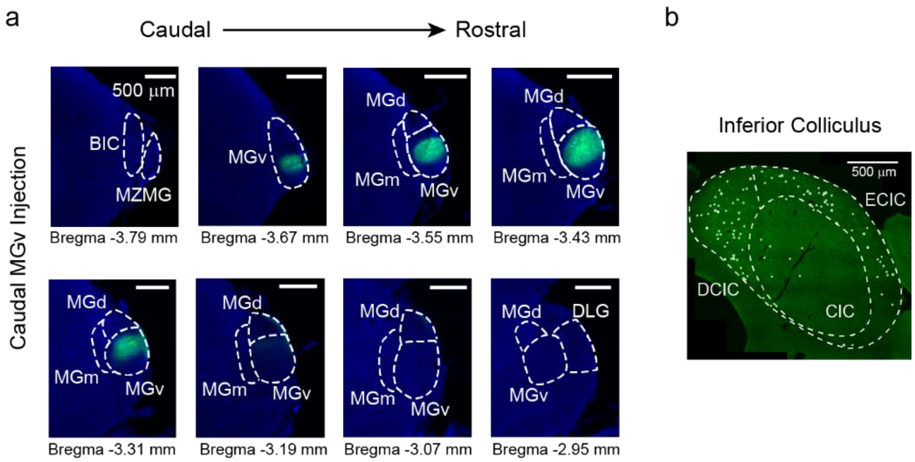


**Supplementary Figure 6. MGN subdivision-targeted tracer injection. (a)** Coronal MGN sections from a representative mouse injected with CTB-488 in cMGv, showing that the tracer was confined to MGv. **(b)** Coronal section of the inferior colliculus in the same mouse, showing input cells (white dots) located predominantly in DCIC and ECIC.

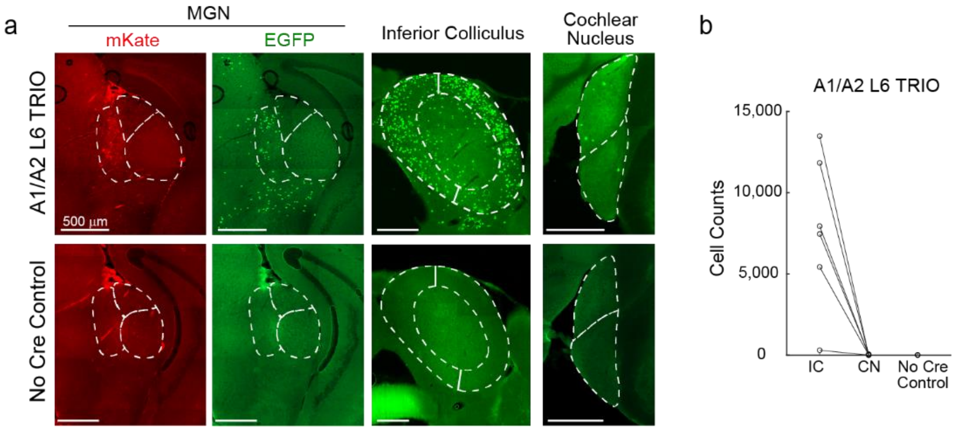


**Supplementary Figure 7. Additional data for disynaptic rabies retrograde tracing. (a)** Coronal brain sections from rabies retrograde tracing. Top: A1 L6 TRIO tracing (MGN and inferior colliculus images are the same as Fig. 5e top). TRIO experiment shows very sparse input cells in the contralateral cochlear nucleus. Bottom: No-Cre control without AAVretro-Cre injection in the cortex. No input cells in no-Cre control. **(b)** Input cell counts in the inferior colliculus (IC) and cochlear nucleus (CN) for TRIO experiments ($n$ = 6 mice including A1 L6 and A2 L6 injections), as well as IC for No-Cre controls ($n$ = 2 mice).

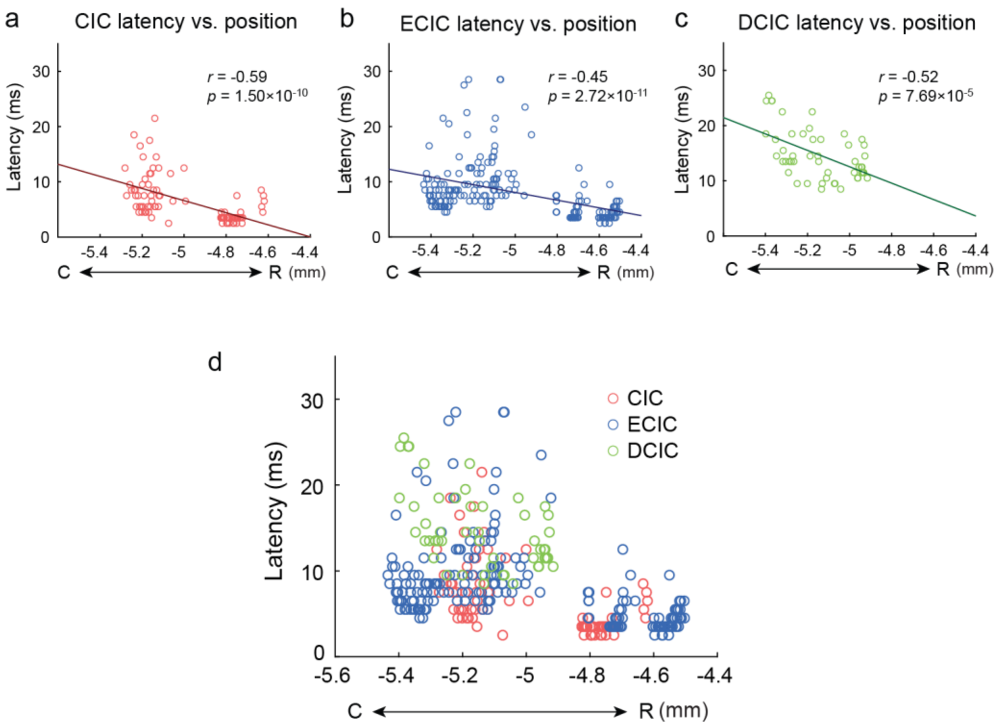


**Supplementary Figure 8. Additional data for inferior colliculus recordings. (a)** Distribution of click response latencies in CIC along the rostrocaudal axis. Red line: linear fit (Pearson's correlation; two-sided t-test). Data same as Fig. 6e. **(b)** Distribution of click response latencies in ECIC along the rostrocaudal axis. Blue line: linear fit. Data same as Fig. 6f. **(c)** Distribution of click response latencies in DCIC along the rostrocaudal axis. Green line: linear fit. **(d)** Overlaid distributions of click response latencies for CIC, ECIC, and DCIC. At the same rostrocaudal coordinates, CIC and ECIC show similar latencies, whereas DCIC latencies tended to be longer.
